# *In Situ* Integration of Porous Polyethylene Nanomembrane with Wound Exudates for Scarless Healing

**DOI:** 10.1101/2024.11.19.624279

**Authors:** Shijie Yuan, Qiao Gu, Xingchen Guo, Xuanxuan Tang, Jiawen Qiu, Linkuo Shang, Julie Qiaojin Lin, Lu-Tao Weng, Ping Gao

**Affiliations:** Multi-functional Polymeric Membranes Research Facility, The Hong Kong University of Science and Technology (Guangzhou), Guangdong, China; Thrust of Advanced Materials, Function Hub, The Hong Kong University of Science and Technology (Guangzhou), Guangdong, China; Green E Materials, The Hong Kong University of Science and Technology (Guangzhou), Guangdong, China; Brain and Intelligence Research Institute, Thrust of Bioscience and Biomedical Engineering, Systems Hub, The Hong Kong University of Science and Technology (Guangzhou), Guangdong, China; Materials Characterization and Preparation Facility (GZ), The Hong Kong University of Science and Technology (Guangzhou), Guangdong, China; Department of Chemical and Biological Engineering, The Hong Kong University of Science and Technology, Hong Kong, China

**Keywords:** Imperceptible wound dressing, pgPE, Self-healing, Scab prevention, Scarless, Breathable, Hypoxia, Follicle regeneration

## Abstract

Achieving scarless wound repair through innovative material designs, such as hydrogels infused with growth factors and cells, has been extensively explored. In this study, we introduce a scarless healing material called pgPE, which is based on a porous and biologically inert ultra-high molecular weight polyethylene (UHMWPE) nanomembrane. This highly flexible nanomembrane conforms closely to the wound site and effectively absorbs wound-associated species into its nanopores, forming pgPE *in situ*. The resulting pgPE creates an ideal environment that fosters physiological metabolism while acting as a barrier against pathogens. Our findings demonstrate that pgPE facilitates complete skin reconstruction, including the regeneration of hair follicles. Additionally, we characterized the immune microenvironment and hypoxic conditions, revealing that pgPE alleviates hypoxia and modulates immune responses, thereby promoting healing towards a scar-free outcome. The integration of wound-associated species within pgPE has been evidenced through three-dimensional Time-of-Flight Secondary Ion Mass Spectrometry (ToF-SIMS) analysis of postmortem samples. This study highlights the potential of personalized solutions that align with physiological systems for enhanced wound healing.

## Introduction

The skin, the largest organ in the human body, is essential for protecting against bacterial invasion and maintaining hydration. Consequently, wound repair—especially the challenge of replacing damaged tissues without fibrosis or scarring—has become a significant research focus ^1–3^. The advancement of new materials is critical for developing effective wound dressings. While traditional polyurethane materials provide effective barriers against bacterial and viral infections, their poor air permeability can hinder metabolic processes essential for healing, as insufficient oxygen supply limits cellular metabolism ^4,5^. On the other hand, hydrogels excel in substance delivery and moisture retention, key features for high-quality wound dressings ^6,7^; however, they too suffer from low air permeability, impeding wound recovery ^8^. A promising direction involves materials that mimic the selective permeability of cell membranes, enabling physiological metabolism around the injury while preventing infections ^9^. This prompts a question: is there a skin-like material that can create an ideal microenvironment for metabolic processes while offering mechanically imperceptible protection against pathogens?

Wound healing is a complex physiological process that involves hemostasis, inflammation, proliferation, and remodeling, with many cells playing crucial roles in regulating healing and scarring, particularly fibroblasts and immune cells at various stages ^10^. Research indicates that appropriately regulating the timing of immune cell involvement and reducing the transformation of myofibroblasts can effectively minimize scarring ^11^. Tissue hypoxia and pathogen invasion threaten healing, which leads to excessive vascularization and unnecessary myofibroblast transformation, potentially resulting in poorly remodel-able granulation and scar tissue ^12^. Therefore, providing a breathable, pathogen-free environment may relieve the need for the activation of the body to recruit immune cells at too early time, reducing myofibroblast transformation or scar tissues. In addition, wound exudate management can also affect wound repair outcomes significantly. Excessive fluidic exudate retention generated from impermeable dressings has been shown to trigger severe inflammatory responses ^13^. On the other hand, heavily dehydrated exudates or scabs resulted commonly from bare wounds have been reported to block air exchange and serve as a breeding ground for bacteria^14–16^. Further, the interaction between the wound and the dressing is another key factor for wound repair. Studies have shown porous materials may facilitate cell adhesion and migration and enhance gas exchange with the external environment for healing enhancement ^17,18^. Additionally, interfacial mechanical interactions at the wound site have been connected closely to the transformation of myofibroblasts ^19^, suggesting the necessity of an imperceptible material for scarless healing. In summary, an effective scar-free dressing should possess skin-like qualities such as breathability, pathogen resistance, humidity regulation, porosity, and mechanical imperceptibility. However, no single material that combines all these properties has been reported so far.

In this study, we provide an innovative solution: a skin-like dressing, called pgPE, which was achieved *in situ* through wound exudates impregnation inside a porous nanomembrane made from ultrahigh molecular weight polyethylene (UHMWPE) ^20^. The nanomembrane possesses porosities greater than 70% and features tunable thicknesses between 20 nm to 1 μm, providing exceptional breathability and ultralow bending stiffness which can conform to the wound site imperceptibly by van der Waals attractions. Three-dimensional (3D) analysis of the nanomembrane detached from the wound using Time-of-Flight Secondary Ion Mass Spectrometry (ToF-SIMS) confirmed the infiltration of electrolytes and amphiphilic wound-associated species into the UHMWPE nanomembrane’s nanopores, implicating a skin-like structure ^21^. This personalized nanofilm transformed *in situ* (pgPE) shows high healing efficacy as demonstrated by macroscopic and histological analyses. An immunostaining analysis revealed that the pgPE maintained physiological metabolisms whilst offering protection against infections. The success of our approach may inform future wound dressing designs to harmoniously integrate with local physiological systems.

## Results

### Design of skin-like pgPE for scarless wound healing

Fig. 1 illustrates our two-step process for creating skin-like pgPE to promote scarless healing. First, we developed self-adhesive porous UHMWPE nanomembrane variants with specific microstructures for pathogen prevention (Fig. 1a). Then, these nanomembranes are converted into pgPE (Fig. 1b) *in situ* through controlled absorption of localized wound exudates into their nanopores. The *in situ* hybridization endowed the pgPE with synergetic properties of the nanomembrane and the natural healing potential of physiological systems.

**Figure 1.**
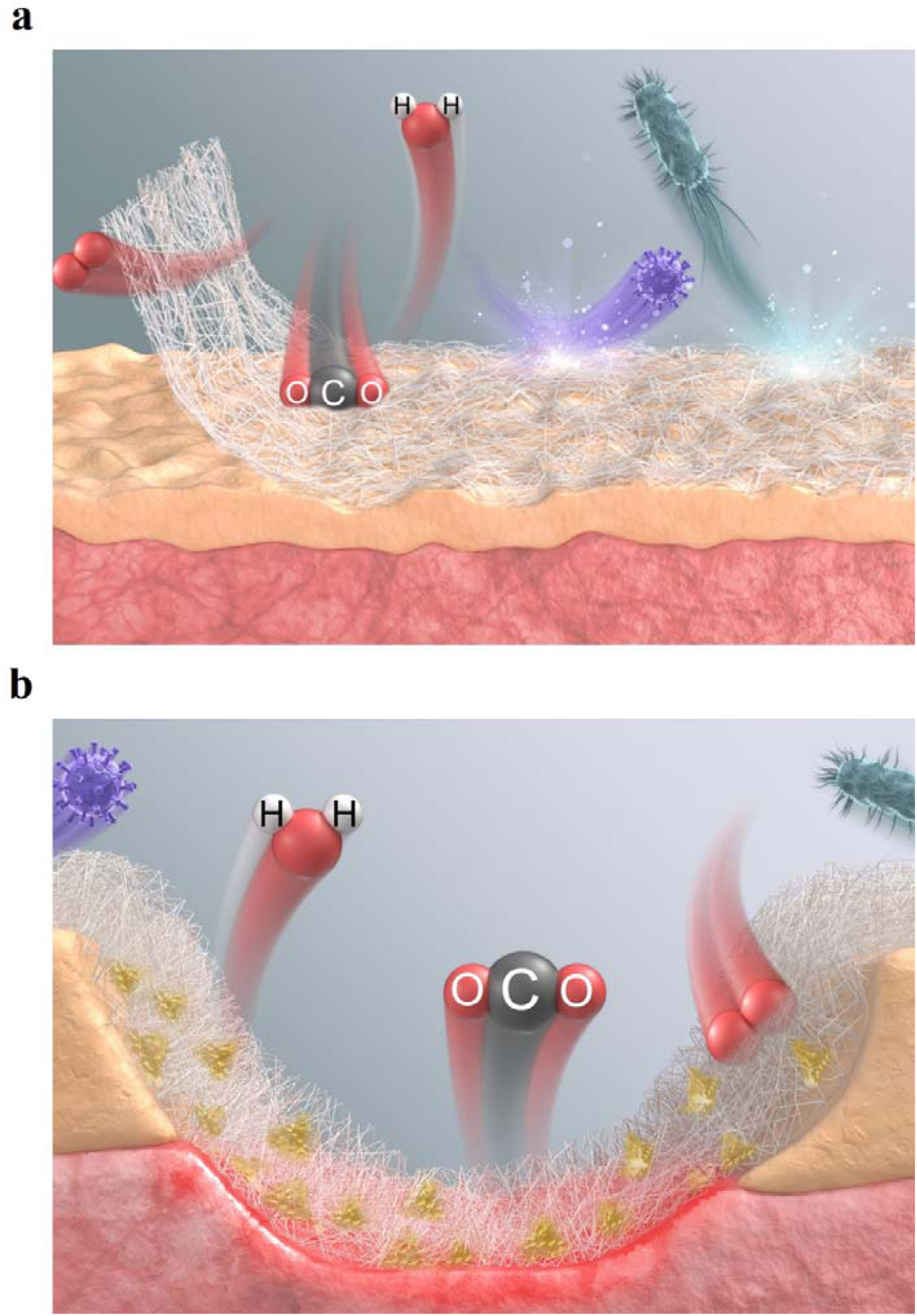
Schematic illustration of our two-step process for creating skin-like pgPE that promotes scar-free wound repair. a) In Step One, a self-adhesive and porous UHMWPE nanomembrane with breathable barrier property is fabricated, as shown by the porous mesh (gray) attached to the skin. b) In Step Two, the nanomembrane formed in Step One is converted into pgPE by incorporating wound exudates after applying it to a 7-mm diameter wound following hemeostasis The fibrous mesh represents the UHMWPE nanomembrane, while the yellow domains represent the locally absorbed wound exudates.

### Fabrication of UHMWPE nanomembranes and *in situ* conversion of pgPE

We first fabricated UHMWPE nanomembranes with tunable microstructures that meet the wound dressing criteria. Figures 2a and 2b (Supplementary Figures 1, and 2) show that these highly transparent nanomembranes are free-standing even at the monolayer level, thanks to their triangulated tessellation of interconnected nanofibrils composed of extended-chain fiber crystals ^22,23^. The material’s high hydrophobicity effectively blocked hydrophilic aerosol particles in the size range of 75 ± 25 nm (Fig. 2c, 2d, Supplementary Fig. 3, 4, 5). But as the source of infections often contains both amphiphilic compounds and hydrophilic species, we created multilayered UHMWPE nanomembranes to ensure full pathogen prevention. Specifically, two variants were fabricated: GPNano-L, with a minimum practical thickness of approx. 80 nm, and GPNano-H, with a maximum self-adhesive thickness of approx. 820 nm, both demonstrating a minimum peel strength of 4.71 N/m (Supplementary Fig. 6). Characterizations using Scanning Electron Microscopy (SEM) and Atomic Force Microscopy (AFM) revealed a multilayered texture that preserved the triangulated topological porous structure observed in the monolayer membrane shown in Fig. 2b (Fig. 2e, 2f). To evaluate their filtration capabilities, we employed *E. coli* and single-stranded Adeno-Associated Virus (ssAAV, diameter = 20-25 nm) as models for permeability testing, confirming that the UHMWPE nanomembranes blocked all bacterial and viral passages (Fig. 2g, Supplementary Fig. 7-10). Gas permeability tests showed that GPNano-L had an air permeability of 5.6×10^6^ L/m²/24h, while GPNano-H demonstrated 7×10^5^ L/m²/24h, making them about 250,000 to 2 million times more permeable than Tegaderm - a commercial dressing (2.74 L/m²/24h) (Fig. 2h) ^24^. Finally, biocompatibility assessments using L929 cells indicated nearly 100% cell viability for the UHMWPE membranes (Supplementary Fig. 11, 12).

**Figure 2.**
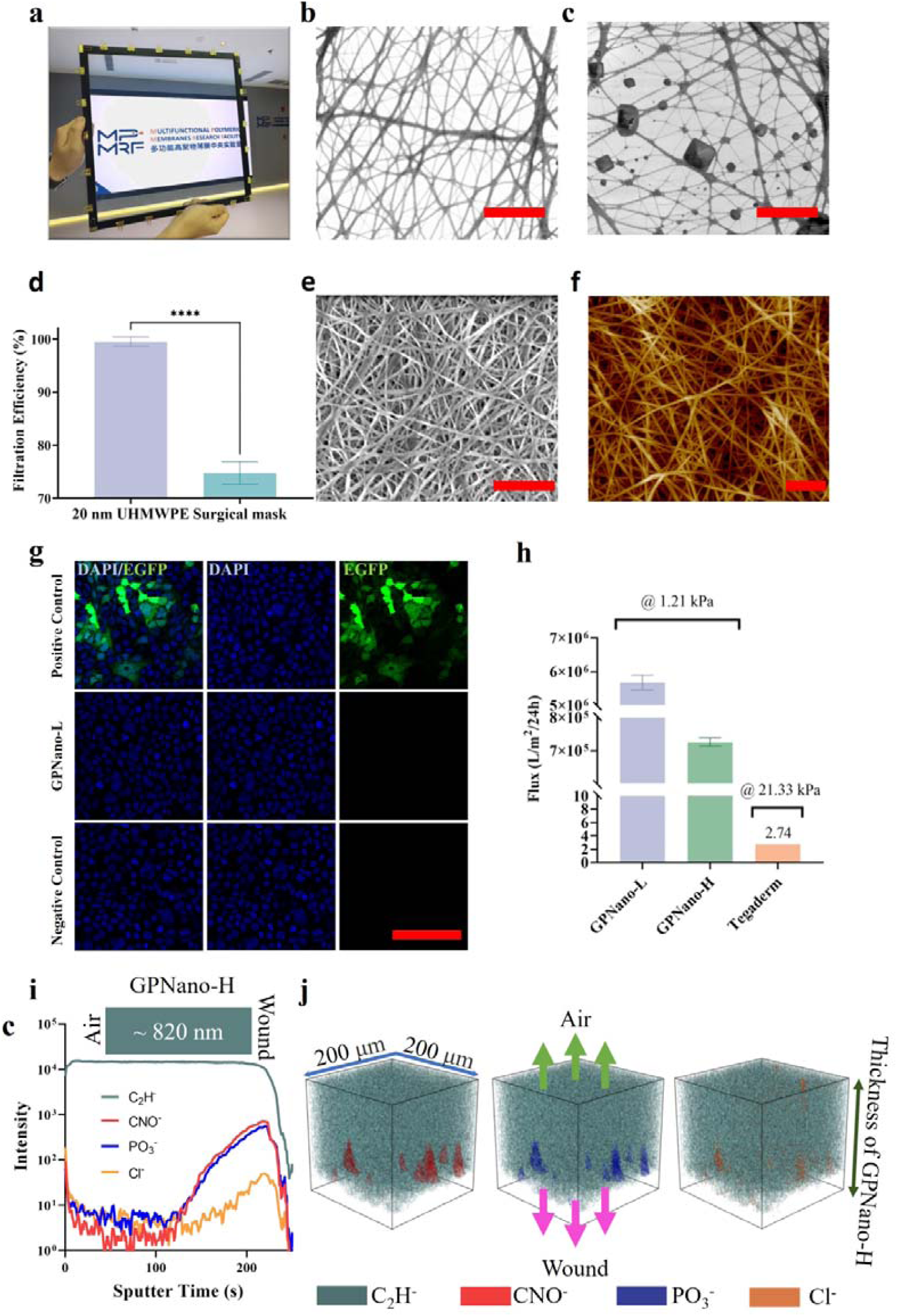
**a)** Photograph showing the free-standing and transparent appearance of the UHMWPE nanomembrane. **b)** Representative SEM image of 20 nm UHMWPE (magnification: 20000x, scale bar: 1μm). **c)** Representative SEM image after the aerosol filtration test (magnification: 20000x, scale bar: 1μm). **d)** Filtration efficiency of 20 nm UHMWPE (n=3, error bar represents the standard deviation). **e)** Representative SEM image of GPNano-H (magnification:50000x, scale bar: 500 nm). **f)** Representative AFM image of GPNano-H (scale bar: 500 nm). **g)** Representative fluorescent images of transfection tests of ssAAV on HEK-293T. Green: transfected cells. **h)** Quantification of gas permeability (n=3 for GPNano-L and GPNano-H, error bar represents the standard error of means). **i)** Negative ToF-SIMS depth profile analysis of GPNano-H (820 nm-thick) from the air site to the wound site. **j)** 3D ToF-SIMS rendering analysis of GPNano-H (the dark green arrow bar=820 nm) in which C_2_H^−^ representing UHMWPE is overlayed with the other ions representing the wound-associated species.

After developing imperceptible UHMWPE nanomembrane variants that meet wound dressing criteria, we converted them into pgPE by integrating localized wound exudates into their nanopores. This conversion was accomplished by applying the UHMWPE nanomembranes to 7 mm-diameter wounds on mice after hemostasis. Integration of wound associated amphiphilic compounds and electrolytes inside pgPE are exhibited in Figure 2i. Due to their exceptional mechanical strength, these nanomembranes can be removed intact from the wound site. We performed a 3D ToF-SIMS analysis of pgPE formed with GPNano-H, which was detached from the wound, at a cryogenic temperature (−120°C), which is necessary to maintain the sample structural integrity and prevent analyte relocation during pumping and analysis ^25^. As shown in Figure 2i, the negative ion intensity of UHMWPE (represented by C_2_H^−^) is uniform throughout the whole GPNano-H film while the ion intensities of various electrolytes (Cl^−^, PO_3_^−^) and amphiphilic chemical species (CNO^−^) associated with the exudates continuously decrease from the surface in contact with the wounding site to the half of the GPNano-H. This suggests that the exudates penetrate the porous UHMWPE, which is fully in agreement with the 3D tomography results (Figure 2j) where the UHMWPE ion (C_2_H^−^) is overlayed with the exudate-associated ions (CNO^−^, PO_3_ ^−^ and Cl^−^). Moreover, the penetration of the exudates can also be viewed from the cross-sectional analysis of CNO^−^, PO_3_^−^ ions (Supplementary Fig 13). Similarly, the infiltration of exudates into porous UHMWPE can be observed in the positive 3D ToF-SIMS analysis of the postmortem samples (Supplementary Figure 14, 15). The low bending stiffness of the UHMWPE nanomembrane, approximately 10^−13^ N.mm ^26^, ensures its conformability, enabling effective integration with exudates, and this integrated structure helps maintain optimal moisture levels. It should b. noted that the GPNano-L, which is about one-tenth of the thickness of the GPNano-H, exhibited some minute scaring, suggesting the need of sufficient thickness for the infiltration of wound exudates. The pgPE formed by the partial filling of exudates in the GPNano-H, in conjunction with its retained wound site conformability enabled it to exhibit a skin-like structure for scar-free healing.

### pgPE facilitates wound healing and reduces scarring

pgPE formed by both variants of UHMWPE nanomembranes demonstrate a strong ability to promote scarless wound healing and accelerate closure rates, as evidenced by macroscopic and microscopic observations along with protein-level characterizations. Figures 3a and 3b (Supplementary Fig. 16) illustrate the progression of wound healing for UHMWPE dressings compared to control and Tegaderm. Clearly, the GPNano-H group showed no visible scabs and scars, while the GPNano-L group had a thin scab and minimal scarring. In contrast, the control group formed prominent, progressively darkening scabs from day 5 to day 9 and clear scarring by day 14. Although no scabs were observed with Tegaderm, its poor air permeability might lead to excessive wound exudates retention, causing prolonged inflammation, and thus impede healing ^27^.

**Figure 3.**
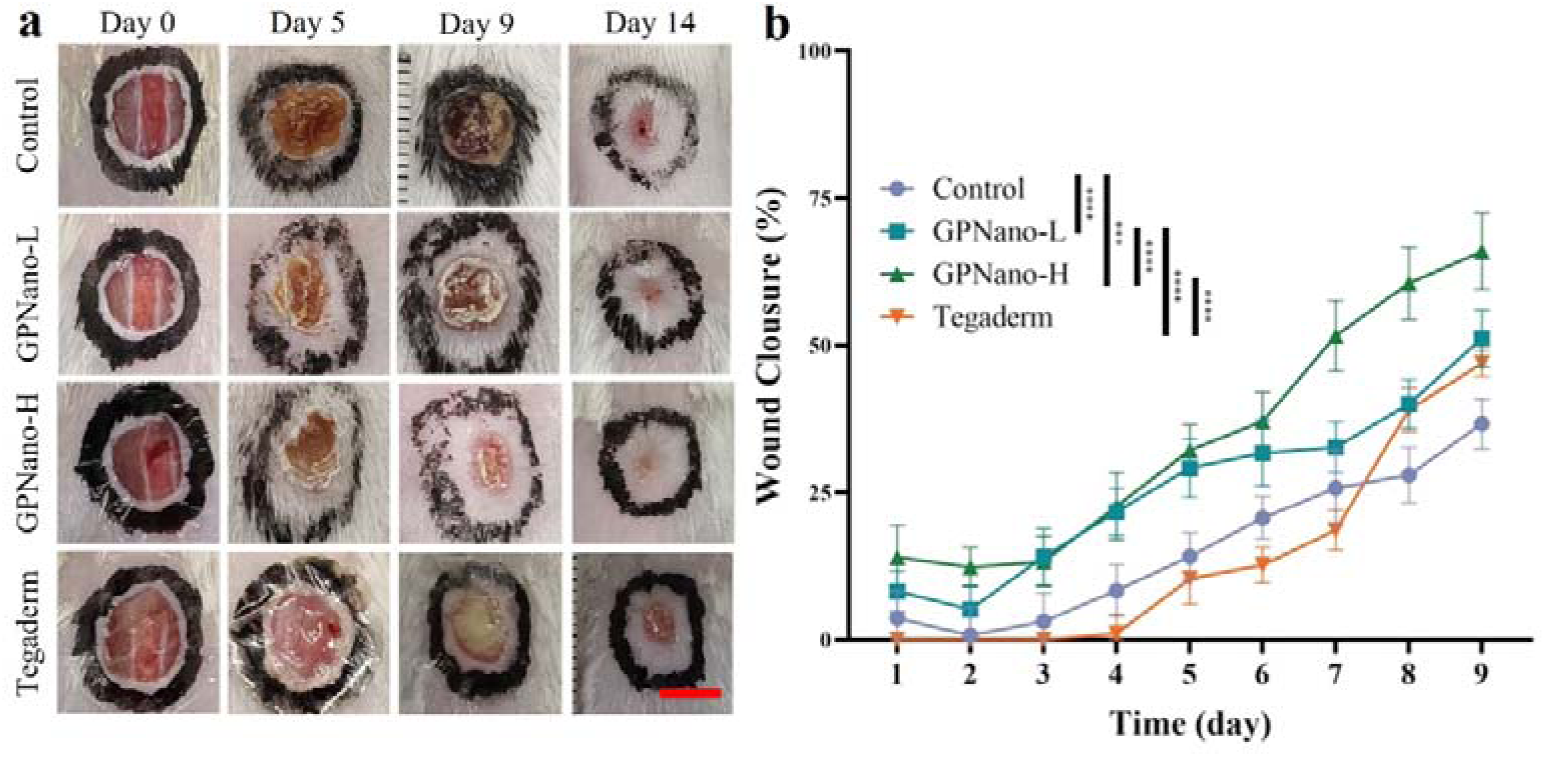
**a)** Representative appearances of wounds on day 0, 5, 9, 14. Scale bar: 7mm. **b)** Quantification of relative wound closure (n=15 (day 1-6), n=10 (day 7-9), two-way ANOVA, *** p<0.0005, **** p<0.0001, error bars represent standard error of means).

The effective healing outcomes induced by pgPE dressings are further corroborated through microscopic investigations. Notably, the GPNano-H group exhibited remarkable healing efficacy, with nearly the entire wound area showing regeneration of hair follicles and sebaceous glands (Fig. 4a). Additionally, the collagen maturity and arrangement (as shown in Masson’s trichrome staining panel) were nearly identical to those found in healthy skin. The GPNano-L group demonstrated slightly inferior performance, consistent with the macroscopic observations. Achieving such a high degree of healing typically requires incorporating additional signaling factors into the dressing to regulate the physiological wound-healing process ^28–31^. Previous studies have indicated that in small full-thickness cutaneous wounds (less than 1 cm × 1 cm) in mice, skin appendages such as sebaceous glands and hair follicles do not regenerate, forming scarring instead ^32,33^. In contrast, the control and Tegaderm groups yielded unsatisfactory results, showing poor appendage regeneration and an immature, disorganized epidermis and dermis. Quantitatively, Fig. 4b-c showcased that both GPNano-L and GPNano-H groups exhibited significantly higher numbers of neo-hair follicles and neo-sebaceous glands. Furthermore, we also quantified the maturity of the wound by measuring the wound area. The results shown in Fig. 4d showed that the GPNano-H group exhibited no immature wound areas, while the GPNano-L group displayed lower levels of immaturity than the control and Tegaderm. Notably, the scarring index of the GPNano-H group was lower than that of the other groups (Fig. 4e). Combined with the in-vivo biocompatibility tests (Supplementary Fig. 17 - 20), these results strongly indicate that the biologically inert UHMWPE nanomembranes facilitate enhanced scarless healing, aligning well with our macroscopic observations.

**Figure 4.**
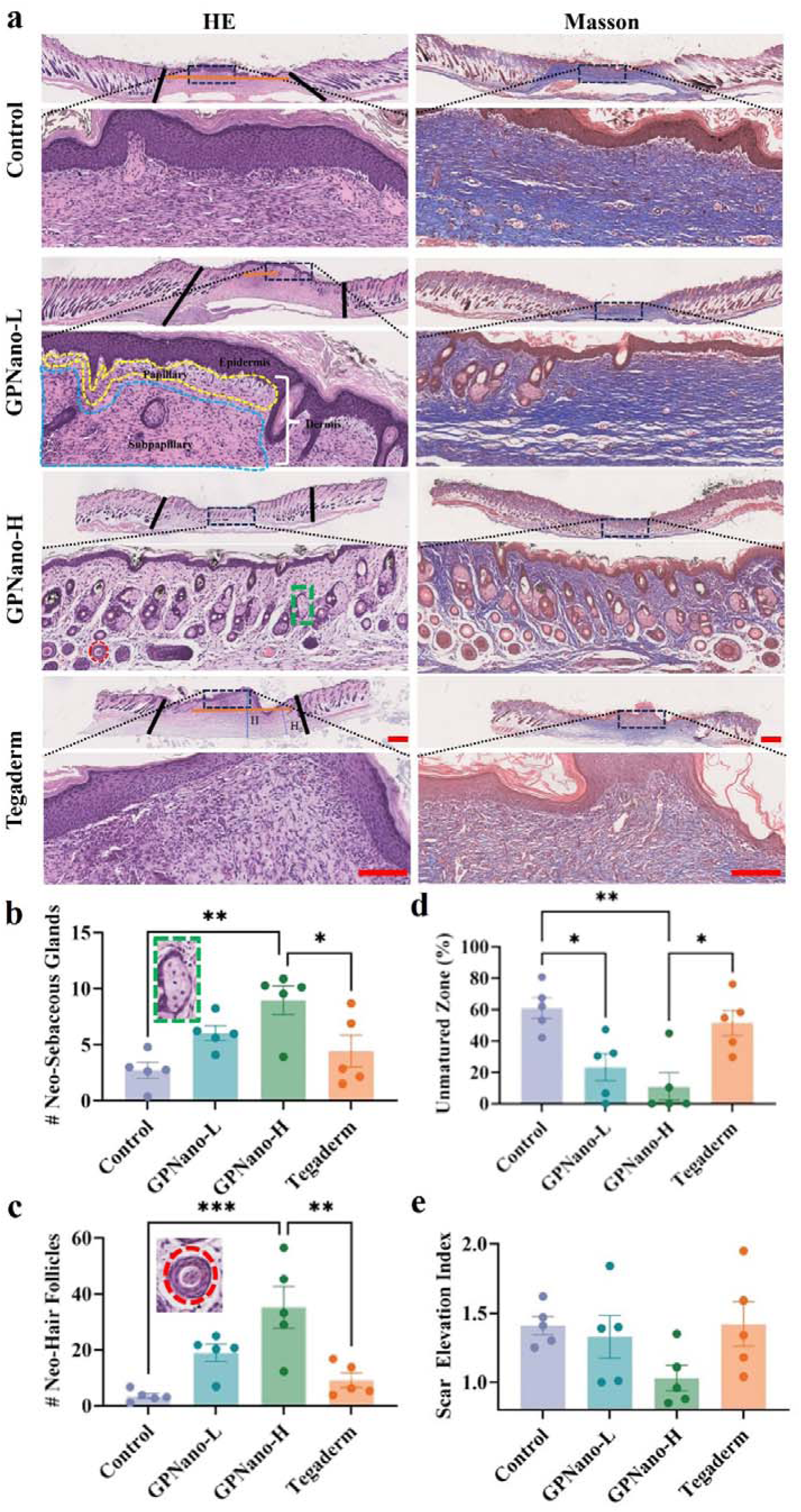
**a)** Representative images of HE-staining and Masson trichrome-staining. (magnification, x25, scale bar: 1 mm; magnification, x200, scale bar: 300 μm). Black lines: wound area. Yellow dashed area: Papillary layer. Blue dashed area: subpapillary layer. Green dashed rectangle: sebaceous gland. Red dashed circle: hair follicle. Orange line: unmatured zone defined by the last two sebaceous glands from the margin of the wound. Blue line and blue dashed line: Height of scar (H) and height of skin (H_0_), respectively. scar elevated index: 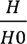. **b-e)** Quantifications analysis of neo-sebaceous glands, neo-hair follicles, unmatured zone, and the Scar Elevation Index. (n=5, one-way ANOVA, *p<0.05, **p<0.001, *** p<0.005, error bars represent the standard error of means).

At the protein level, the UHMWPE nanomembrane groups demonstrated their scarless healing capability by exhibiting significantly lower expression levels of α-smooth muscle actin (α-SMA) (Fig. 5a, 5b) compared to the control and Tegaderm groups. Previous studies suggest that α-SMA is predominantly expressed in myofibroblasts and vascular endothelial cells, showing a strong correlation with scar formation through wound fibrosis ^34–36^.

**Figure 5.**
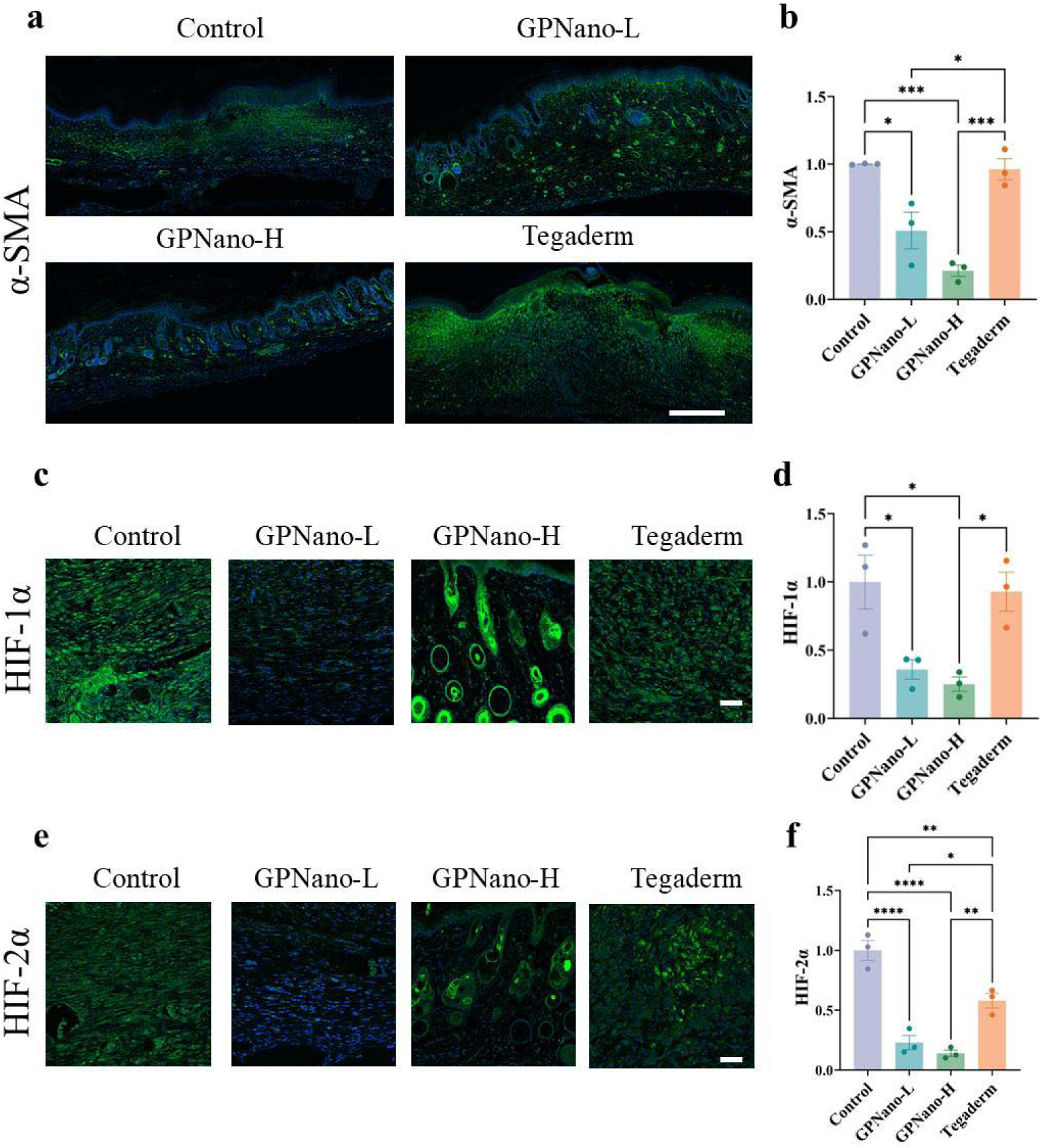
**a)** Representative immunofluorescence staining of α-SMA. (Green: α-SMA. Blue: DAPI Scale bar: 500 μm). **b)** Quantification analysis of the calibrated total expression (IntDen) of α-SMA. (n=3, one-way ANOVA. *p<0.05, **p<0.001, *** p<0.0005, **** p<0.0001, error bars represent the standard error of means). **c**) Representative immunofluorescence staining of HIF-1α. (Green: HIF-1α, blue: DAPI, Scale bar: 50 μm). **d)** Quantification analysis of the calibrated total expression (IntDen) of HIF-1α per cell (n=3, one-way ANOVA. *p<0.05, **p<0.001, *** p<0.0005, **** p<0.0001, error bars represent the standard error of means). **e**) Representative immunofluorescence staining of HIF-2α. (Green: HIF-1α, blue: DAPI, Scale bar: 50 μm). **f)** Quantification analysis of the calibrated total expression (IntDen) of HIF-2α per cell (n=3, one-way ANOVA. *p<0.05, **p<0.001, *** p<0.0005, **** p<0.0001, error bars represent the standard error of means).

Having confirmed that the artificial skin characteristics of pgPE exhibit wound healing efficacy and minimize scarring at macroscopic, microscopic, and protein levels, we will further explore the key factors contributing to these healing outcomes.

### pgPE regulates the wound microenvironment to enhance natural metabolism

The skin-like pgPEs exhibit their effectiveness by regulating the wound microenvironment, promoting enhanced natural metabolism. This regulation is evidenced by reduced hypoxic levels and diminished inflammatory responses.

As sufficient oxygen exchange is essential for wound healing, we first assessed the severity of hypoxic induction in the wounded tissues when covered with pgPE by immunohistochemistry of hypoxia-inducible factor 1-alpha (HIF-1α), a well-studied hypoxia indicator. HIF-1α was found at low levels in both the GPNano-L and GPNano-H groups, whereas its expression levels were significantly higher in both the control and the Tegaderm group (Fig. 5c, 5d). The elevated HIF-1α levels in the control and Tegaderm groups can be attributed to their low air permeability; Tegaderm’s limited air permeability was illustrated in Fig. 2h, and the scabs formed in the control group hindered air exchange (Fig. 3a). A similar trend was observed for hypoxia-inducible factor 2-alpha HIF-2α (Fig. 5e, 5f). Thus, the high air permeability of pgPE helps alleviate hypoxic conditions in the wound, facilitating better oxygen transmission.

Various immune factors and inflammatory cells are considered next. Previous studies have shown that M2 macrophages can reduce inflammation and promote follicle regeneration ^37^. Their presence in the wound area can be visualized through immunofluorescence staining of CD206, a specific surface antigen of M2 macrophages. Figures 6a and 6b show that the pgPEs exhibited at least 50% higher number of CD206^+^ cells in the wound compared to both the control and Tegaderm groups. This implicates that the moisture management property provided by the skin-like pgPEs fosters the transformation of M2 macrophages, thereby enhancing wound healing and collagen reconstruction, which helps prevent scar formation. We also examined the expression levels of IL-4, which has been identified as a risk factor for scar formation due to its role in fibroblast proliferation and collagen density ^38,39^. From days 5 to 14, we found that IL-4 levels in the pgPEs were consistently lower than those in the Tegaderm and control groups, even in regions with similar immaturity (Fig. 6c, 6d, Supplementary Fig. 21). The expression profile of IL-13 mirrored that of IL-4, with the pgPEs showing reduced levels of IL-13 as well (Supplementary Fig. 22). This suggests that the rapid and effective healing observed in the pgPEs does not coincide with an increase in IL-4 and IL-13 expression, thereby reducing the risk of wound fibrosis and scar formation, consistent with our earlier observations (Fig. 3a).

**Figure 6.**
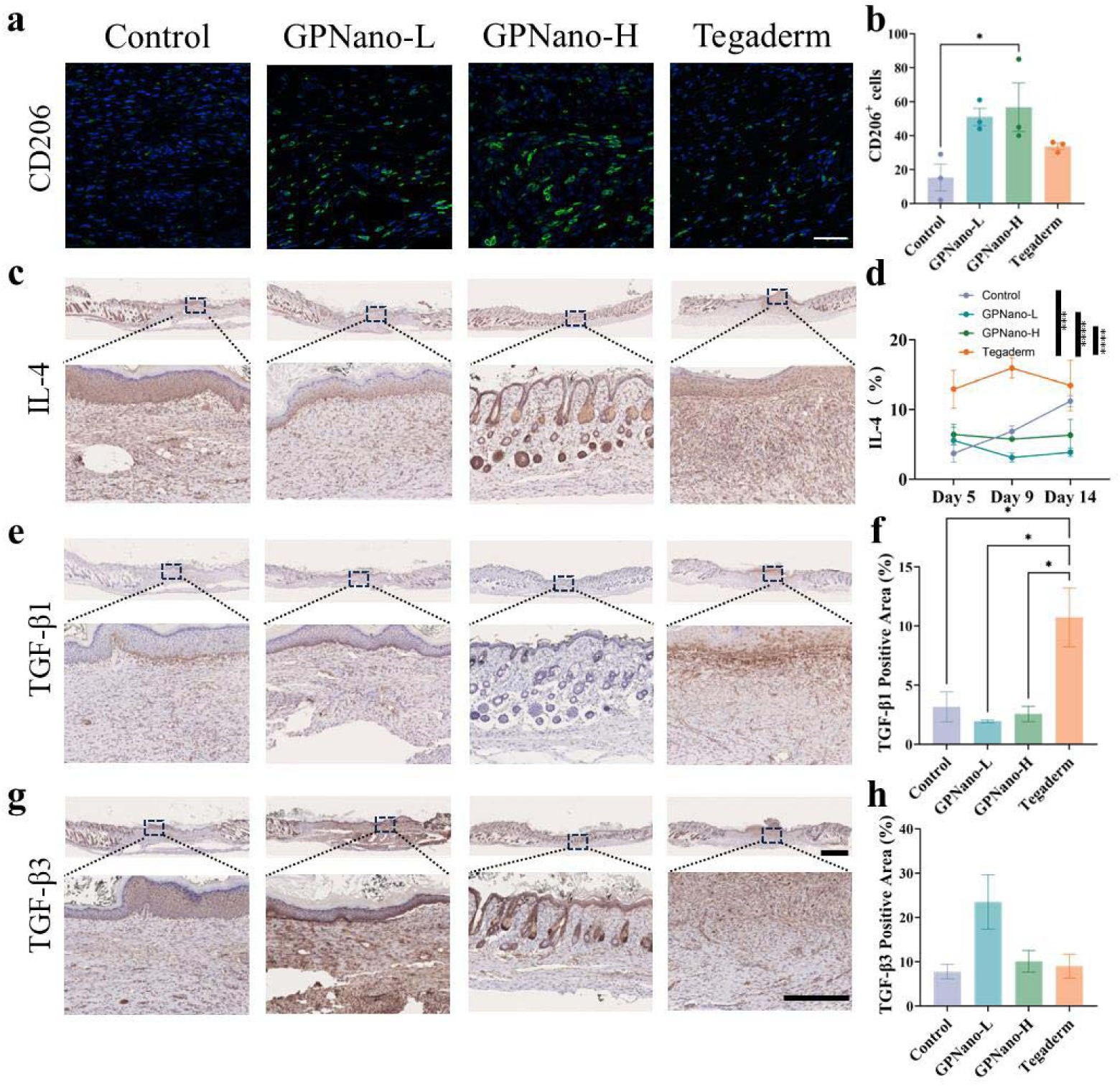
**a)** Representative immunofluorescence staining of CD206. (Green: CD206^+^ cells. Blue: DAPI Scale bar: 50 μm). **b)** Quantification analysis of CD206^+^ cells per view. (n=3, one-way ANOVA. *p<0.05, error bars represent the standard error of means). **c)** Representative immunohistochemistry staining of IL-4 on day 14. (Brown: IL-4. magnification, x25, scale bar: 1 mm; magnification, x200, scale bar: 300 μm). **d)** Quantification analysis of IL-4 expression from day 5 to day 14. (n=3, one-way ANOVA. **p<0.01, error bars represent the standard error of means). **e)** Representative immunohistochemistry staining of TGF-β1. (Brown: TGF-β1. magnification, x25, scale bar: 1 mm; magnification, x200, scale bar: 300 μm). **f)** Quantification analysis of TGF-β1 expression. (n=3, one-way ANOVA. **p<0.01, error bars represent the standard error of means). **e)** Representative immunohistochemistry staining of TGF-β3. (Brown: TGF-β3. magnification, x25, scale bar: 1 mm; magnification, x200, scale bar: 300 μm). **f)** Quantification analysis of TGF-β3 expression. (n=3, one-way ANOVA. Error bars represent the standard error of means).

Based on these findings, we hypothesize that the changes in the downstream pathways caused by the inflammatory microenvironment are key to achieving faster healing and scarless wound repair. Transformative growth factor (TGF)-β1 is a well-known downstream cytokine linked to wound healing and scar formation ^40,41^. Although TGF-β3 also promotes cell proliferation, it is not typically associated with scarring. In wound healing, the ratio of TGF-β3 to TGF-β1 is often used as a characteristic to assess scaring risk ^42,43^. Our immunohistochemical analysis revealed comparable TGF-β1 expression levels between the pgPE formed by GPNano-L and control groups (Fig. 6e). In the pgPE formed by the GPNano-H group, where complete healing was almost reached, TGF-β1 levels were barely detectable, whereas the Tegaderm group exhibited significantly elevated TGF-β1 levels (Fig. 6f). Conversely, TGF-β3 expression was widespread in the GPNano-L group, low in the well-healed GPNano-H group, and minimal in the control and Tegaderm groups (Fig. 6g, 6h). Quantitative analysis revealed that the GPNano-L group had a significantly higher TGF-β3/TGF-β1 ratio compared to the other groups (Supplementary Fig. 23). Notably, the expression levels and ratios in the GPNano-H group were not surprising, as the group was nearing complete healing and had entered the remodeling phase, moving beyond the primary TGF-regulated growth phase.

### Outlook

We have demonstrated a skin-like platform, pgPE, for scar-free wound repair, which forms *in situ* through the integration of a nanomembrane with localized electrolytes and amphiphilic species from exudates, thus creating an ideal environment for effective skin reconstruction. The absence of scabs, low levels of hypoxia (HIF-1α), extremely low α-SMA, and a significantly high CD206 indicate a naturally metabolic environment akin to healthy skin, facilitating humidity regulation, air exchange, mechanical imperceptibility for inflammation regulation, and pathogen protection. The successful full regeneration of skin without scarring underscores the synergistic interplay between physiological systems and the UHMWPE nanomembrane dressing. This research may also inform future designs of wound dressing that require the desirable integration with physiological systems.

## Acknowledgments

We would like to express our gratitude to the MCPF(GZ) of HKUST(GZ) for their invaluable technical support and guidance. We also thank Guangdong HWT for their professional oversight of animal ethics and protocols. Additionally, we appreciate the technical assistance and support provided by Guangdong Solidgood Limited and Fumengdao Guangzhou Limited. We also appreciate the King’s Flair International (Holdings) Limited. We also thank Prof. Zhuoyi Liang for providing the HEK-293T cell line and Yaxiong Fang for technical support. This work is supported by the Tsinghua Zhujiang Research Institute of Pearl River Delta (to P.G.) and the Guangzhou Science and Technology Program City-University Joint Funding Project (Project No. 2023A03J0001) (to J.Q.L.).

## Contributions

S.Y. and P.G. designed the materials and research and wrote the manuscript. S.Y. conducted most experiments without further indications. L-T. W. and J.Q.L. provided significant suggestions during manuscript preparation. Q.G. performed the aerosol permeability test and SEM. X.G. performed the physical characterizations. X.T. performed biocompatibility tests and bacterial permeability tests. J.Q. performed the ToF-SIMS. L.S. performed the virus permeability tests. All authors reviewed the manuscript and offered feedback.

## Methods

### 1. Materials

UHMWPE nanomembranes were developed and fabricated in our group, with slight modifications ^44^. Briefly, a low-entanglement-density gel film was biaxially stretched by BRUCKNER KARO 5.0 in a semi-solid state at 90-130°C. The draw ratio of GPNano-H is 100-200 times, while that of GPNano-L is 400-600 times. The stretched gel films were constrained and extracted to remove the oil. The extraction system consisted of HPLC-grade n-hexane, ethyl acetate, ethanol, and water. After extraction, the gel films were vacuum-heated to remove water and other solvent residues for 72 hours. Finally, the samples were sealed in a positive-pressure room and disinfected for 24 hours using ultraviolet light.

A detailed list of the commercially available materials utilized in this study, including supplier names, catalog numbers, and CAS numbers, is available in Supplementary Data 1. All primers were purchased from Sangon Biotech.

### 2. Characterization methods

#### (a) Peel Strength test

A UHMWPE sample was cut into a 2 cm x 2 cm piece and was immediately applied to the subject’s forearm by van der Waals attraction facilitated by alcohol spraying^26^. A 90° peel strength test was conducted using an Instron 68TM, with a peeling speed of 50 mm/s. The other piece was applied to adjacent skin after the test was completed, and the same tensile test was performed. Detailed schematic diagrams and images can be found in Supplementary Fig. 24.

#### (b) Aerosol Permeability Test

The aerosol permeability of the monolayer UHMWPE (∼20 nm) was performed on TSI 8130A.

#### (c) Gas Permeability Test

We tested UHMWPE samples on the HBYIQI HB-TQY-01 with a pressure drop of 1.21 kPa and measured the time it took for 100 mL of gas to pass.

#### (d) Scanning Electron Microscopy

SEM micrographs of the thinnest UHMWPE were obtained using a hot field emission SEM instrument (JEOL 7800F) with an acceleration voltage of 2 kV. The sample film was prepared by suspending UHMWPE on a perforated bare transmission electron microscope copper grid with 200 mesh. No conductive coating was applied to the film.

The SEM micrographs of GPNano-H and GPNano-L samples were obtained using a cold field emission SEM instrument (Hitachi Regulus 8230) with an acceleration voltage of 1 kV. The GPNano samples were put on a silicon wafer and then coated with a platinum thin film using Quorum Q150T.

#### (e) Atomic Force Microscopy

AFM images of GPNano samples were obtained using a Bruker Dimension Icon instrument with a Scanasyst-Air probe in PeakForce mode. Samples were attached to a surface of an atomically flat mica sheet.

#### (f) Time-of-Flight Secondary Ion Mass Spectrometry

ToF-SIMS 3D analyses were carried out on an IONTOF M6 hybrid SIMS instrument (Ion-ToF GmbH, MuCnster, Germany). This instrument is equipped with a 30 keV bismuth liquid-metal ion source for analysis and an argon cluster ion source for sputtering. A focused 10 keV Ar ^+^ beam scanning over an area of 500 μm × 500 μm was used to sputter through the detached UHMWPE nanofilms. Sample imaging was performed over an area of 200 μm × 200 μm at the center of the sputter region using a 30 keV Bi_3_ ^+^ bunched cluster beam. The target pulsed current of the Bi_3_ ^+^ beam was ∼0.3 pA. Each imaging scan contained 256 × 256 pixels. The whole profiling analysis was run in the non-interlaced mode, consisting of cycles of short pulses of Bi_3_^+^primary beam for analysis followed by long pulses of Ar_1302_^+^ beam for sputtering. Charge compensation was realized using a low-energy flood gun (∼21 eV). All SIMS spectra, chemical mapping, and 3D reconstruction were analyzed using SurfaceLab 7.3

To preserve structural integrity and avoid analyte relocation during pumping and analysis under ultra-high vacuum ^25^, GPNano-H samples were prepared and analyzed in the following way. Once detached from the wounds, the GPNano-H samples (∼ 1.4 cm * 1.4 cm) were laid down on a piece of silicon wafer, with the 4 edges fixed using SEM tape in order to keep the samples as flat as possible. Samples were pre-cooled in the Loadlock chamber at −120°C for ∼ 2 hours under dry N_2_ and then pumped down. Once the Loadlock chamber reached a vacuum of ∼10^−6^ mbar, they were transferred to the main chamber, in which the sample stage was kept also-120°C. All the SIMS analyses were performed at −120°C and a pressure of ∼8.9 X 10^−9^ mbar.

### 3. Biological tests

#### (a) Virus Permeability Test

2.4 mL of ultrapure water was added to a 35 mm cell culture dish, and the surface was sealed with GPNano-L, ensuring that the upper surface of the membrane was in a relaxed state. Then, 0.6 mL of ultrapure water was added to apply pressure, creating a separation of only one layer of membrane between the two layers of ultrapure water. Next, 1 µL of ssAAV suspension, with a total of 10^9^ GC, was added to the topside ultrapure water. The dish was placed at 4°C for 24 hours. After 24 hours, the upper liquid and GPNano-L were carefully discarded, and the lower aqueous phase was preserved for viral gene extraction, according to TIANamp Virus DNA/RNA Kit (TIANGEN, DP315).

We used a 20 µL PCR reaction system, which included 2 µL of extracted viral genomic DNA, 0.6 µL each of forward and reverse primers, 10 µL of 2X Rapid Taq Master Mix (Vazyme, P222), and 6.8 µL of nuclease-free ultrapure water. First, the mixture was pre-denatured at 95°C for 5 minutes, followed by 30 amplification cycles, each consisting of 15 seconds of denaturation at 95°C, 30 seconds of annealing at 57°C, and 30 seconds of extension at 72°C, with a final hold at 4°C. After PCR completion, the PCR products were detected by 2% agarose nucleic acid gel electrophoresis. The sequences of the GFP primers are as follows: F: GGTGAACTTCAAGATCCGCC; R: CTTGTACAGCTCGTCCATGC.

#### (b) Transfection Test

HEK-293T cells were seeded into a glass-bottom 6-well plate. When the confluence reached 30%, the upper culture medium was discarded, fresh medium was added, and GPNano-L was applied as described above. To the clear water above the GPNano, 10 µL of 10^13^ GC/µL ssAAV virus suspension was added, and the plate was incubated at 37°C for 72 hours. Subsequently, confocal microscopy imaging and cell RNA extraction were performed. Confocal imaging was performed by AX Confocal Microscopy (NIKON).

After washing the cells twice with PBS, 500 µL of Trizol (ThermoFisher, 15596026CN) was added to each well of a 6-well plate and incubated at room temperature for 5 minutes. Then, 100 µL of chloroform was added, and the mixture was vigorously shaken and incubated at room temperature for 3 minutes. After centrifugation, the upper aqueous phase was collected into a new RNase-free centrifuge tube. The RNA was washed sequentially with isopropanol and 75% ethanol, discarding the supernatant each time. The tube was left to dry at room temperature for 5 minutes, then DEPC water was added to dissolve the RNA, and the RNA concentration was measured for subsequent rtPCR and qPCR.

The reverse transcription program was completed according to the protocol of the PrimeScript™ RT Reagent Kit (Takara, RR047A), with an RNA input amount of 0.5 micrograms per sample. Each sample was pre-diluted to ensure consistent concentration. Subsequently, real-time quantitative PCR was performed using the TB Green Premix Ex Taq II Kit (Takara, RR820A). The primer sequences used were as follows:

GFP-F: GGTGAACTTCAAGATCCGCC;

GFP-R: CTTGTACAGCTCGTCCATGC;

GAPDH-F: ACGACCACTTTGTCAAGCTCATTTC;

GAPDH-R: GCAGTGAGGGTCTCTCTCTTCCTCT.

#### (c) *In-vitro* biocompatibility test

Direct contact test: L929 cells were seeded into a 96-well plate and cultured until reaching 80% confluency. A 1 cm * 1 cm of GPNano-L, GPNano-H, or nothing was added into different wells, with each material replicated three times. After 24 hours, samples were removed, and the cell viability of each group was tested according to the Cell Counting Kit-8 (Biosharp, BS350A).

Leachate test: Cells were prepared as above and after, supernatants were replaced by serum-free medium. In a sterile environment, GPNano-L and GPNano-H were cut into 7 cm x 7 cm pieces and soaked in 10 mL of serum at room temperature for 48 hours. Then, they were added to the corresponding wells at a 10% volume ratio, with normal serum used as a negative control. The remaining CCK-8 tests were performed as described above.

#### (d) Animal Care

All animal experiments and procedures were approved by the ethics committee, with approval number SYXK(Yue) 2023-0249. BALB/c male mice, aged 6-8 weeks and weighing 18-22 g, were purchased from the Guangdong Medical Laboratory Animal Center (SCXK (Yue) 2022-0002)). All animals were acclimated for one week before the experiments. The animals were housed under standard conditions (temperature 23 ± 2°C, humidity 55 ± 5%), with a 12-hour light/dark cycle, and had free access to food and drinking water ad libitum. After surgery, animals were housed individually.

#### (e) Mouse Full-Thickness Cutaneous Wound Model

We employed a full-thickness cutaneous model in mice to demonstrate the healing efficacy of the two UHMWPE variants and the potential for natural healing. The surgery was conducted under sterile conditions, with mice anesthetized using isoflurane. Throughout the surgical process and during recovery from anesthesia, a heating pad was used to maintain body temperature. The dorsal skin was depilated using hair removal cream, followed by careful shaving of fine hairs with a razor, and then disinfected with povidone-iodine. Sterile punches (7 mm in diameter) were used to create a full-thickness excisional wound in the center of the back.

The experimental groups are as follows (n=5): 1) Control group - 5 days; 2) GPNano-L group - 5 days; 3) GPNano-H group - 5 days; 4) Tegaderm group - 5 days; 5) Control group - 9 days; 6) GPNano-L group - 9 days; 7) GPNano-H group - 9 days; 8) Tegaderm group - 9 days; 9) Control group - 14 days; 10) GPNano-L group – 14 days; 11) GPNano-H group - 14 days; 12) Tegaderm group - 14 days;

After grouping, different materials were applied, and the mice were allowed to recover from anesthesia. Daily photographs of the wound were taken and quantitatively analyzed by ImageJ. The materials were inspected twice daily for adherence, with reapplication as needed. At each time point (5, 9, and 14 days), animals were sacrificed by isoflurane anesthesia followed by the collection of blood from the retinal venous plexus for anticoagulation, and cervical dislocation. Skin tissue, heart, liver, spleen, and kidneys were collected. Except for the skin, the remaining tissues were fixed in 10% neutral formalin. The skin tissue was divided along the spinal axis into two halves: one half was preserved in liquid nitrogen, and the other half was fixed in 10% polyformaldehyde.

#### (f) Histological Analysis

Tissues were embedded in paraffin, sectioned into 4 µm slices, and stained with hematoxylin and eosin (H&E) as well as Masson’s trichrome stain. All slides were scanned using a NanoZoomer S60 (Hamamatsu) and observed with NDP.view. The wound analysis was performed by three researchers in a double-blind manner. The wound area was defined by residual muscle tissue, and hair follicles and sebaceous glands within the wound area were counted by three researchers, with the average value taken. The immature region was determined by the distance between the most distant mature hair follicles at both ends of the wound, while the scar elevation index was calculated as the ratio of the maximum wound thickness to the thickness at the wound edges.

#### (g) Immunobiological Chemistry (IHC)/ Immunofluorescence (IF) Analysis

The sections were processed by AS330 PLUS Automatic IHC (Dartmon). In brief, sections were dewaxed in xylene, rehydrated in a gradient, subjected to citrate buffer antigen retrieval, and blocked with 3% bovine serum albumin in 10% serum. For IHC, sections were incubated with H_2_O_2_ blocking before serum, followed by primary antibodies (IL-4 Abcam ab9811, 1:100; IL-13 Invitrogen PA5-102574, 1:100; TGF-β1 Abcam ab215715, 1:100; TGF-β3 Abcam ab15537, 1:100), secondary antibody (Dartmon 51457), and diaminobenzidine (Dartmon 51457). Sections were scanned and analyzed using the same instrument and software in the histological analysis. For IF, sections were incubated with primary antibodies (α-SMA Abclonal A7248 1:50; HIF-1α Abcam ab228649, 1:200; HIF-2α Invitrogen PA1-16510, 1:100; CD206 Abclonal A21014, 1:50), secondary antibody (Goat Anti Rabbit AF488 Abcam ab150077, 1:200), DAPI, and observed by AX Confocal Microscope (NIKON).

#### (h) Statistical Analysis

Statistical analysis was performed with GraphPad Prism 10. Each figure legend contains corresponding analysis methods. In brief, data are shown as the mean ±[standard error of means. One-way ANOVA was used to analyze the single variable multiple groups data and two-way ANOVA was used to analyze the multiple variables data.

