## Supplementary Figures for "*In Situ* Integration of Porous Polyethylene Nanomembrane with Wound Exudates for Scarless Healing"

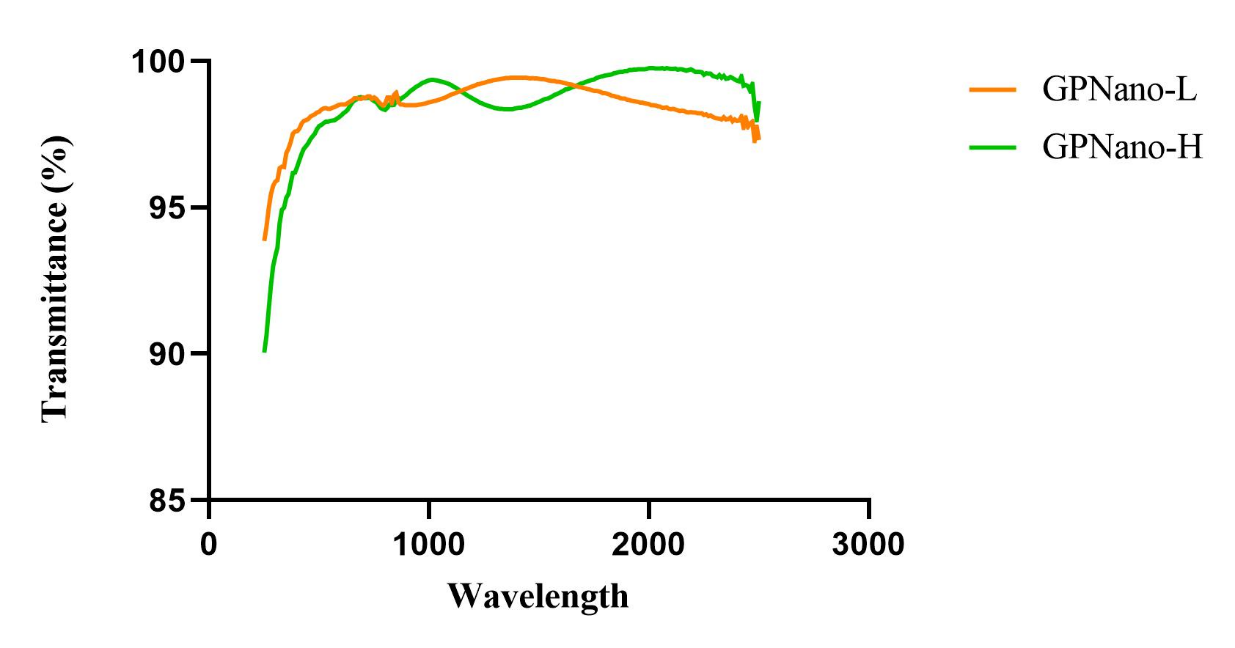

**Supplementary Fig. 1** UV-Visible spectra of two UHMWPE variants, measured by LAMBDA 1050+ (Perkin Elmer).

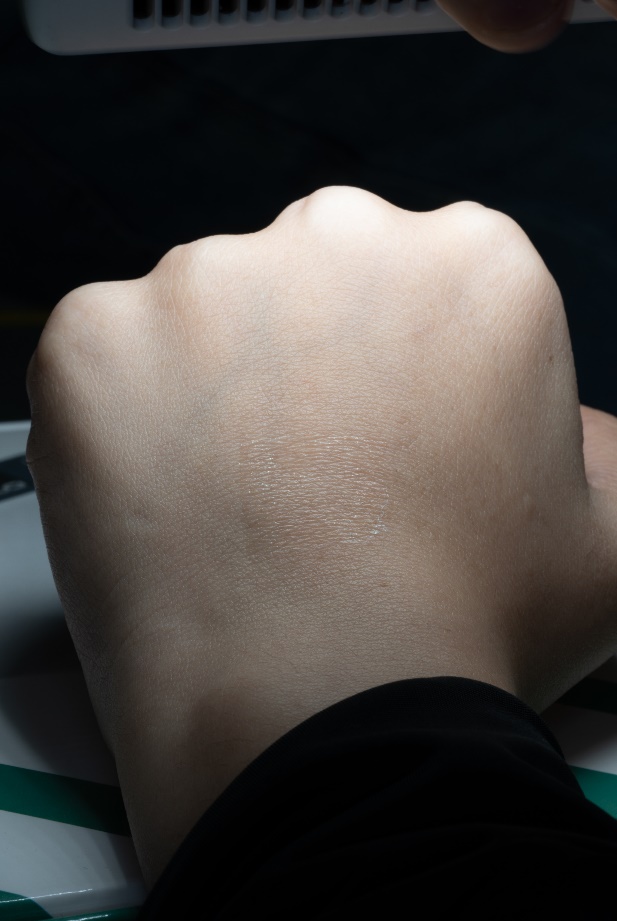

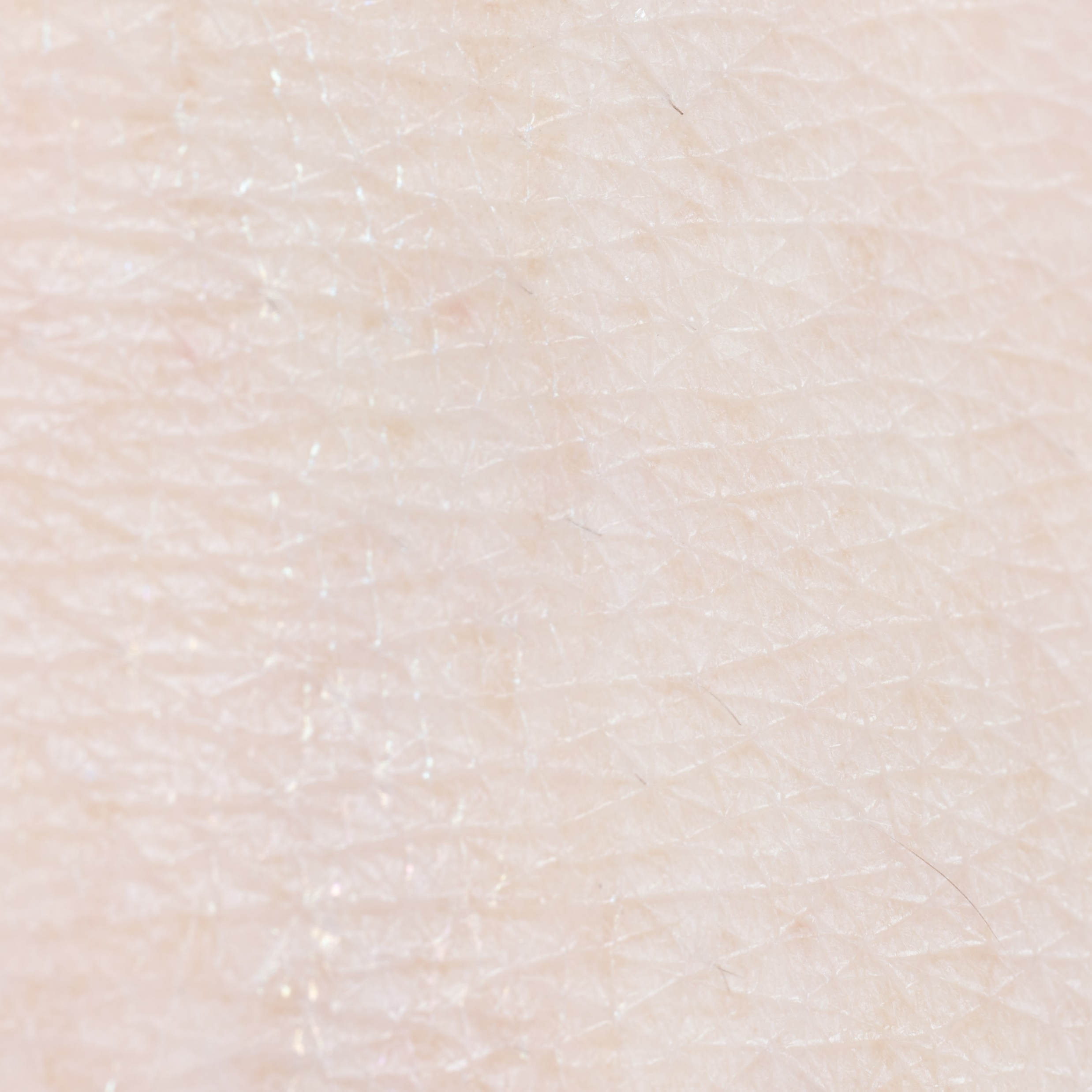

GPNano-L

GPNano-L

**Supplementary Fig. 2** Appearance of GPNano-L on hand (red dashed lines: margins of GPNano-L).

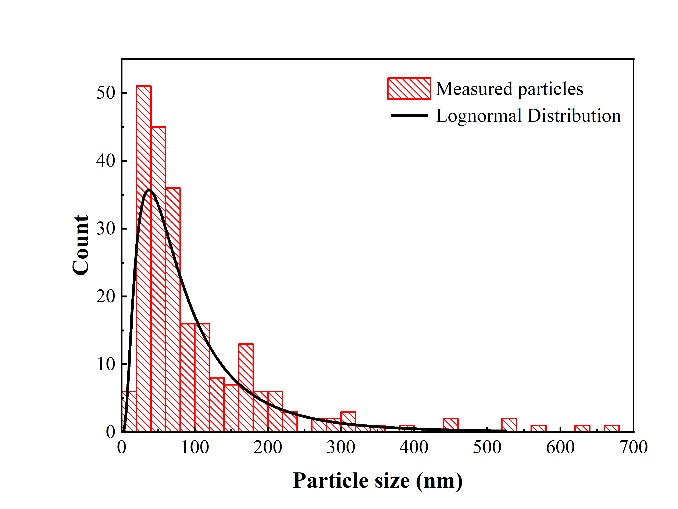
**Supplementary Fig. 3** Schematic illustration of aerosol filtration test and particle size distribution of aerosols.

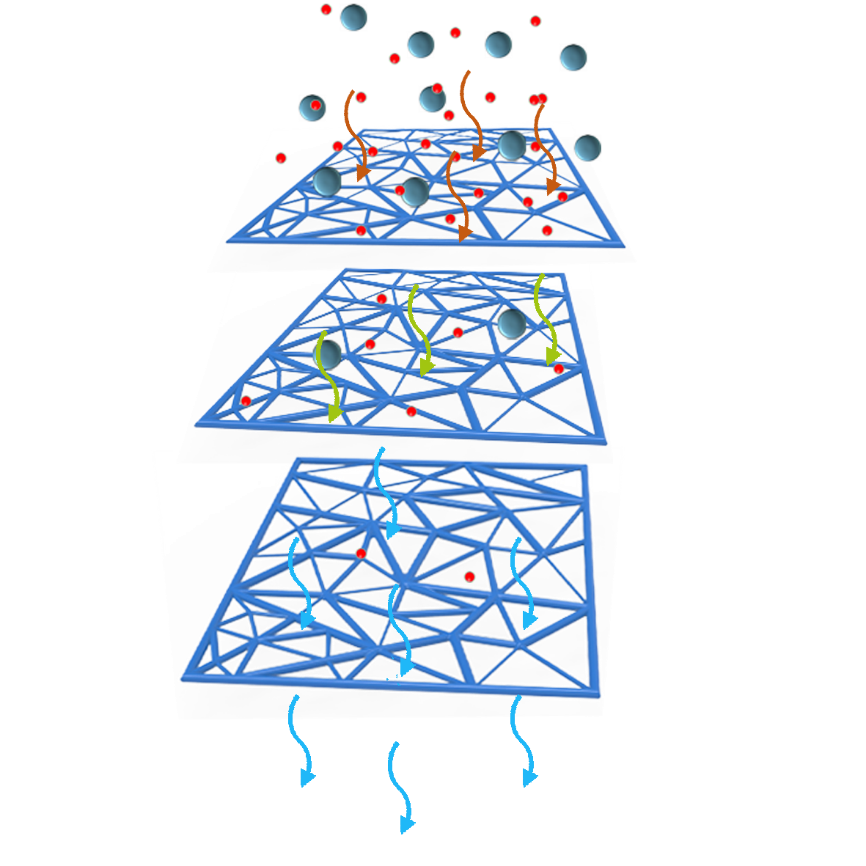

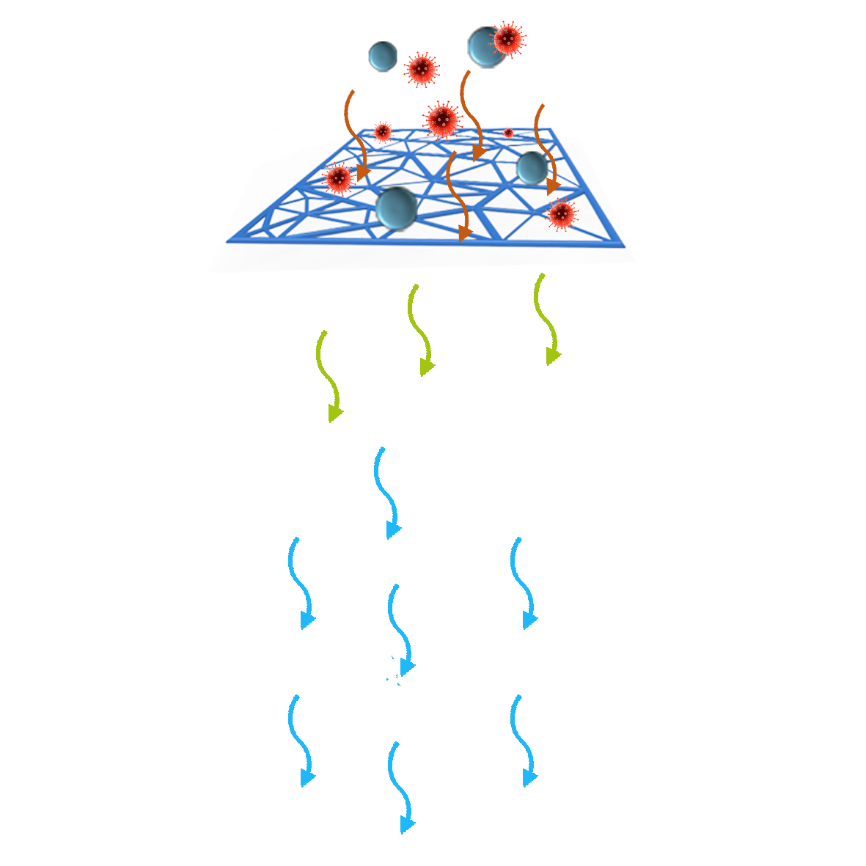

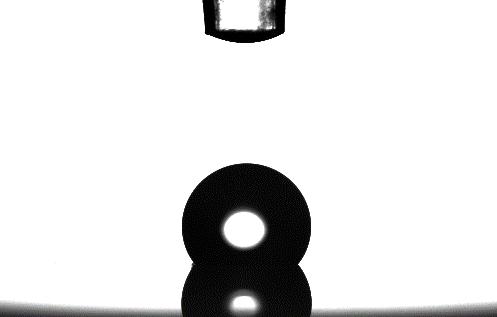

$$CA\approx130^{\circ}$$

**Supplementary Fig. 4** Representative contact angle of GPNano-L.

Pore Size Distribution-Histogram

Pore Size Percentage

| Pore Size (nm) | 2023-08-24 MEMBRANE 03 |
| --- | --- |
| 20.55 to 21.00 | 0.189618 |
| 21.00 to 22.00 | 0.320159 |
| 22.00 to 23.00 | 0.760638 |
| 23.00 to 24.00 | 1.32633 |
| 24.00 to 25.00 | 3.52725 |
| 25.00 to 26.00 | 4.57066 |
| 26.00 to 27.00 | 5.80719 |
| 27.00 to 28.00 | 34.2261 |
| 28.00 to 29.00 | 27.8949 |
| 29.00 to 30.00 | 3.332 |
| 30.00 to 31.00 | 2.35935 |
| 31.00 to 32.00 | 1.36504 |
| 32.00 to 33.00 | 0.699824 |
| 33.00 to 34.00 | 0.675099 |
| 34.00 to 35.00 | 0.752971 |
| 35.00 to 36.00 | 1.52528 |
| 36.00 to 36.76 | 2.00784 |

**Supplementary Fig. 5** Pore size distribution data of GPNano-L. (average pore size = 27.8 nm)

**Supplementary Fig.**
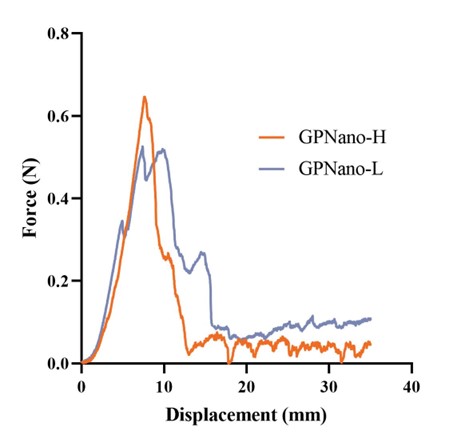
**6** Peel strength measurements of different UHMWPE variants.

**Supplementary Fig. 7** Representative images of *E. coli* permeability test. Colonies represent the existence of *E. coli*.

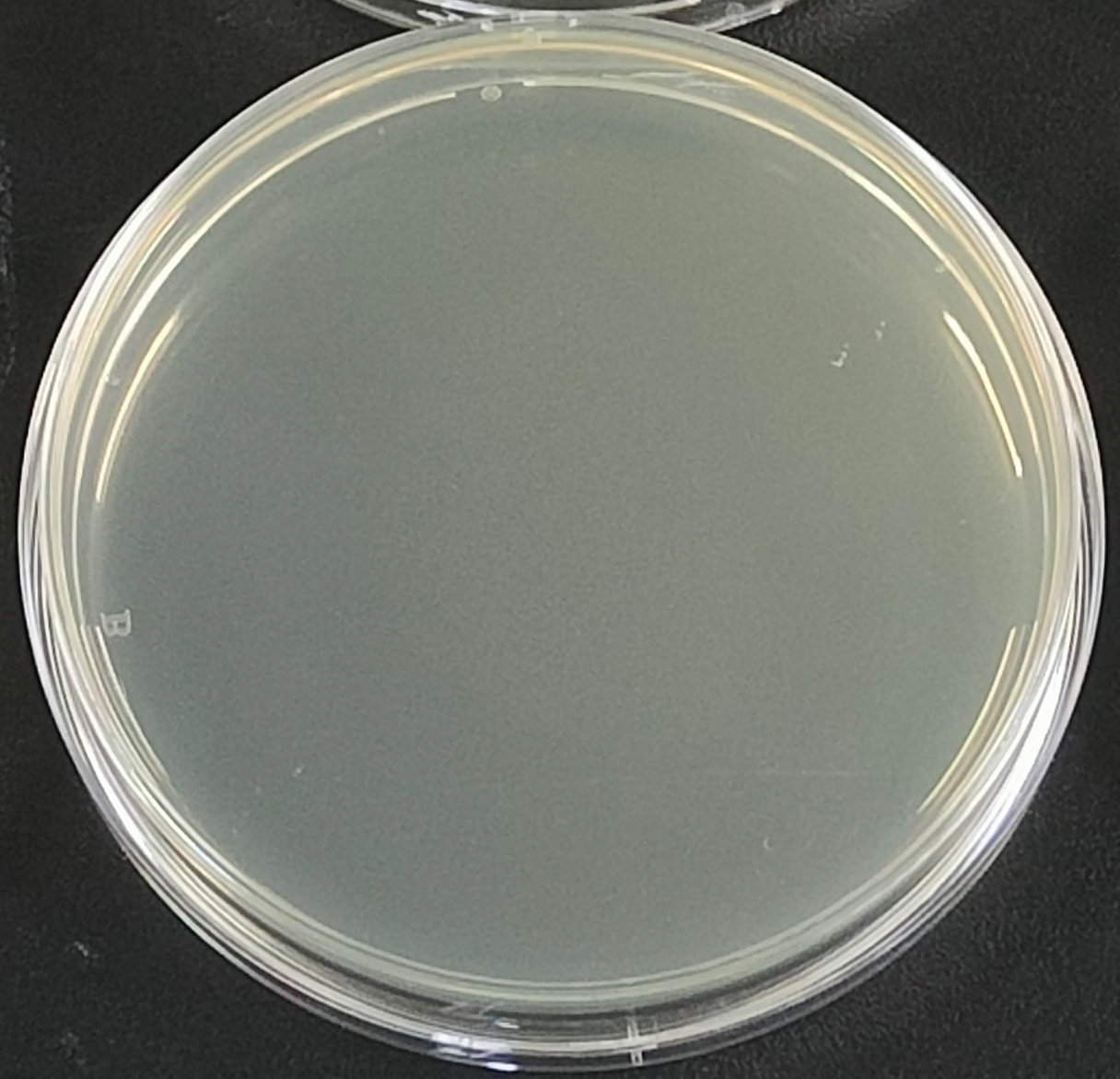

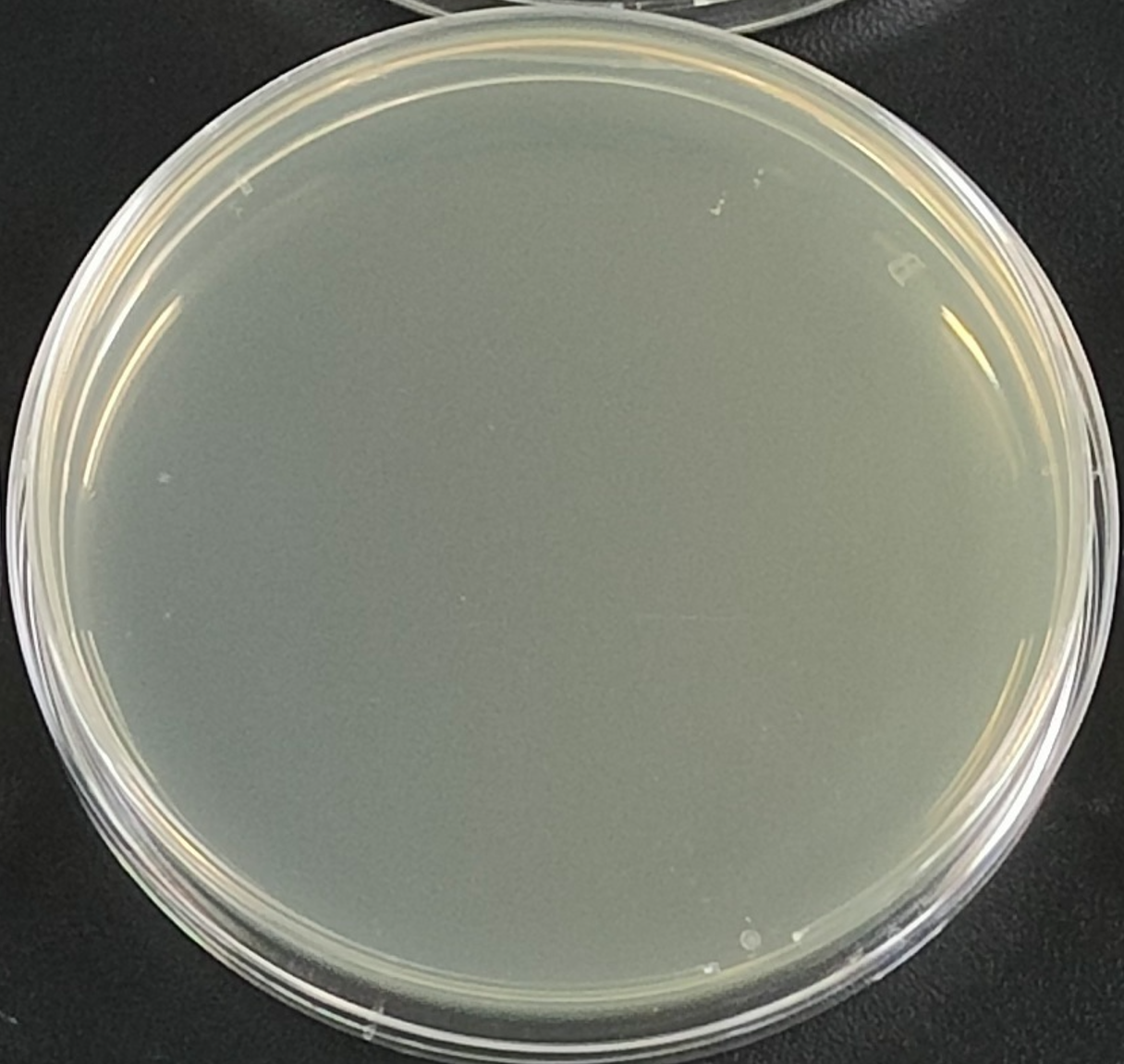

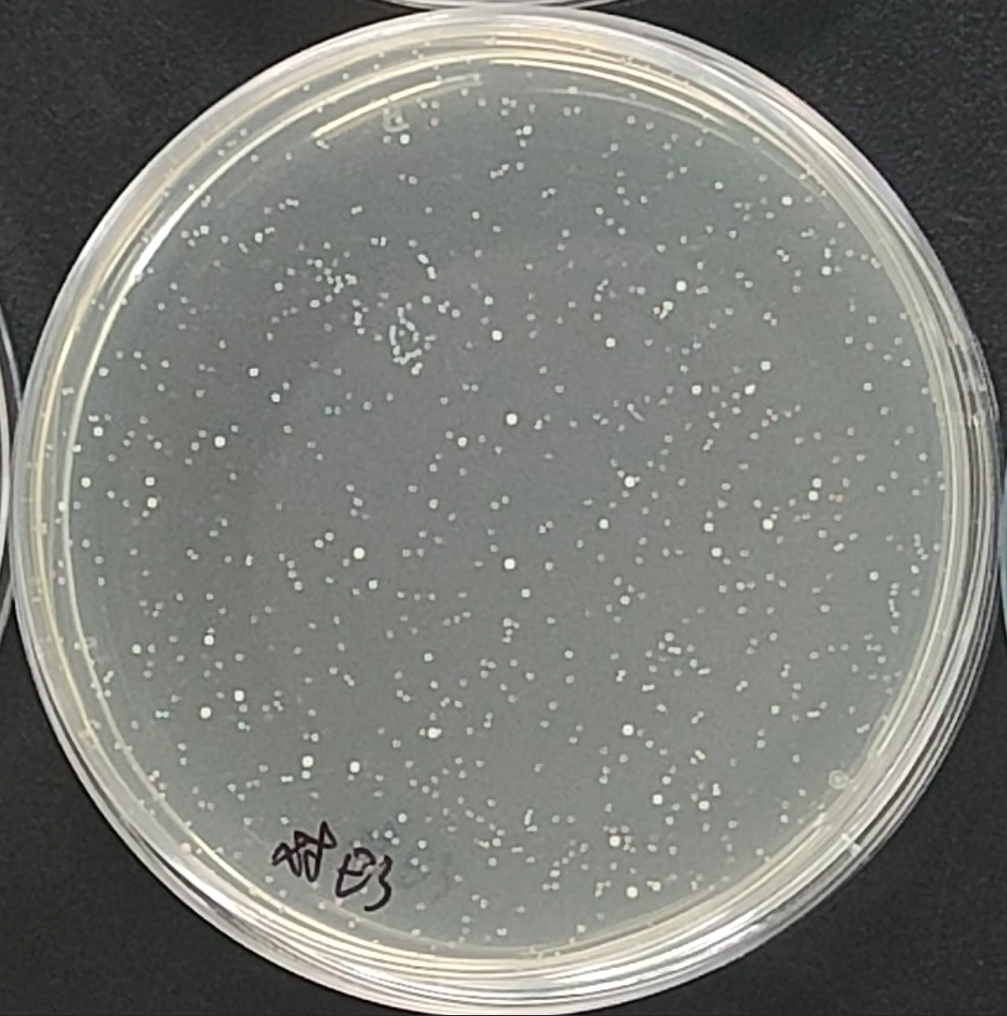

Control

GPNano-H

GPNano-L

Marker

Pure Water + GPNano-L #1

Pure Water + ssAAV #1

Pure Water + ssAAV #2

Pure Water + ssAAV #3

Pure Water + ssAAV + GPNano-L #2

Pure Water + ssAAV + GPNano-L #1

Pure Water + ssAAV + GPNano-L #3

Pure Water + GPNano-L #2

Pure Water + GPNano-L #3

100 bp

200 bp

300 bp

400 bp

500 bp

600 bp

700 bp

800 bp

900 bp

1000 bp

1500 bp

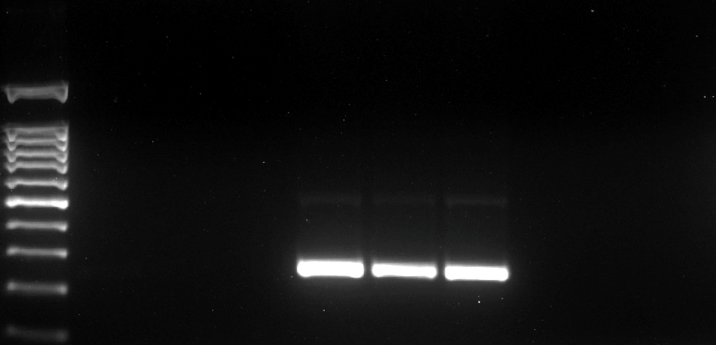

**Supplementary Fig. 8** Representative PCR image of ssAAV permeability test.

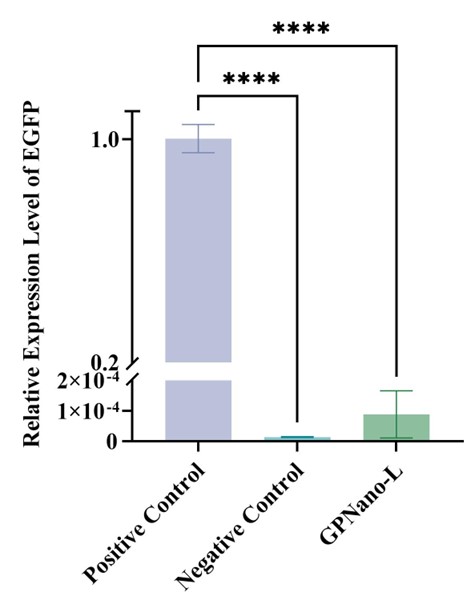

**Supplementary Fig. 9** Quantification of ssAAV transfection test on HEK-293T measured by qPCR (n=3, error bar represents the standard error of means, ****p<0.0001).

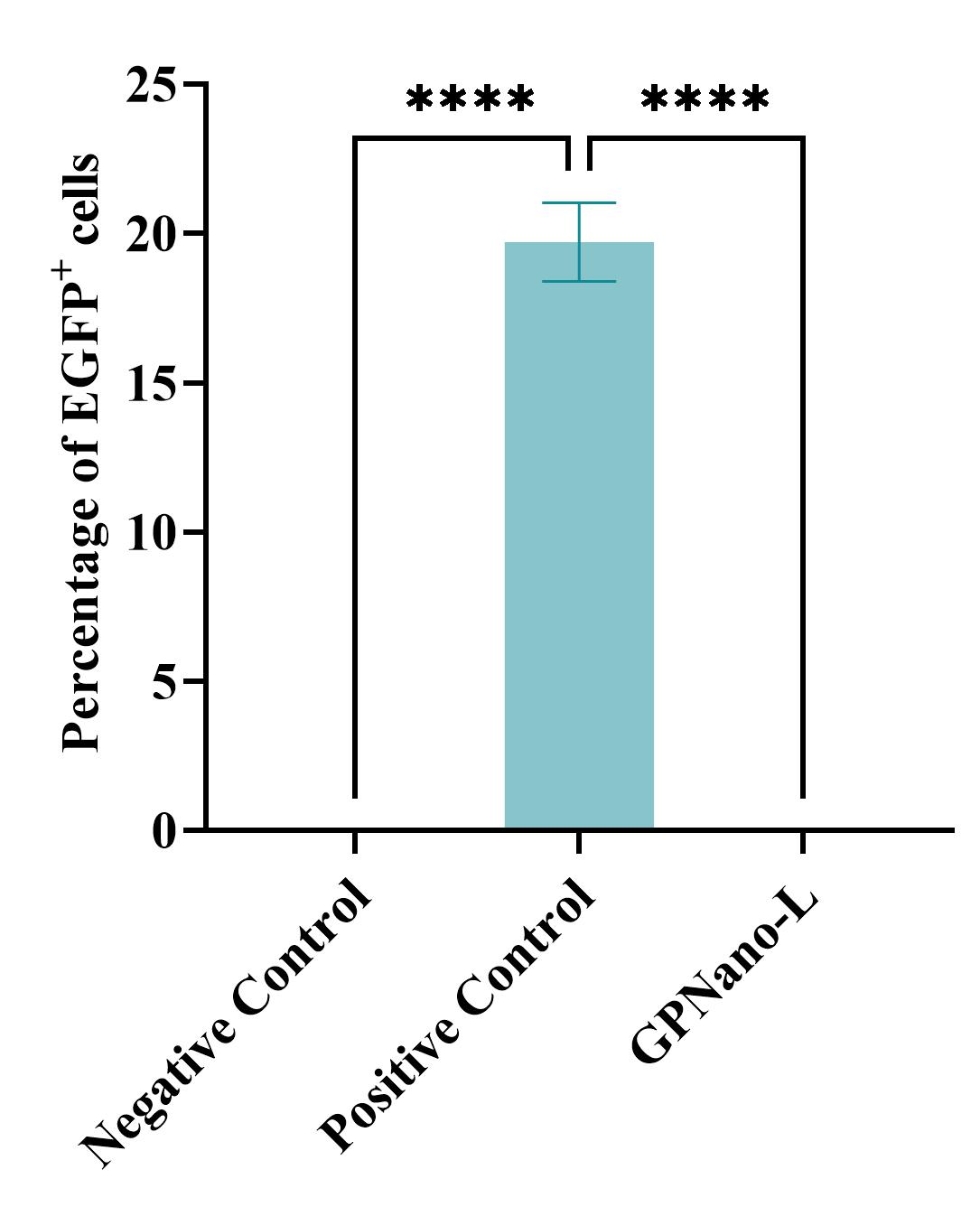

**Supplementary Fig. 10** Quantification of ssAAV transfected cells (EGFP^+^) percentage (n=6, error bar represents the standard error of means, ****p<0.0001).

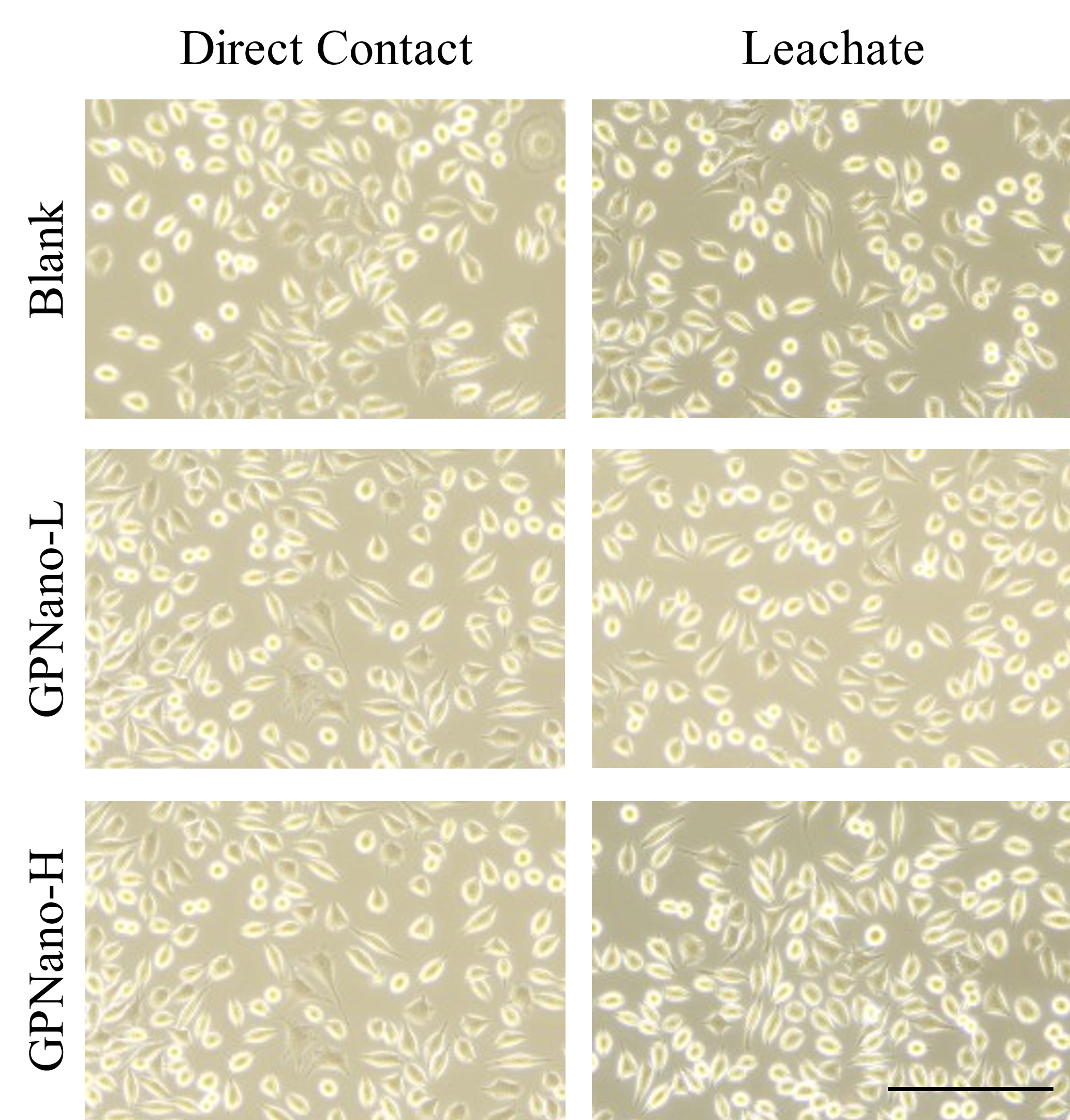

**Supplementary Fig. 11** Representative images of the L929 biocompatibility test. (Magnification: 20x; scale bar: 200 μm)

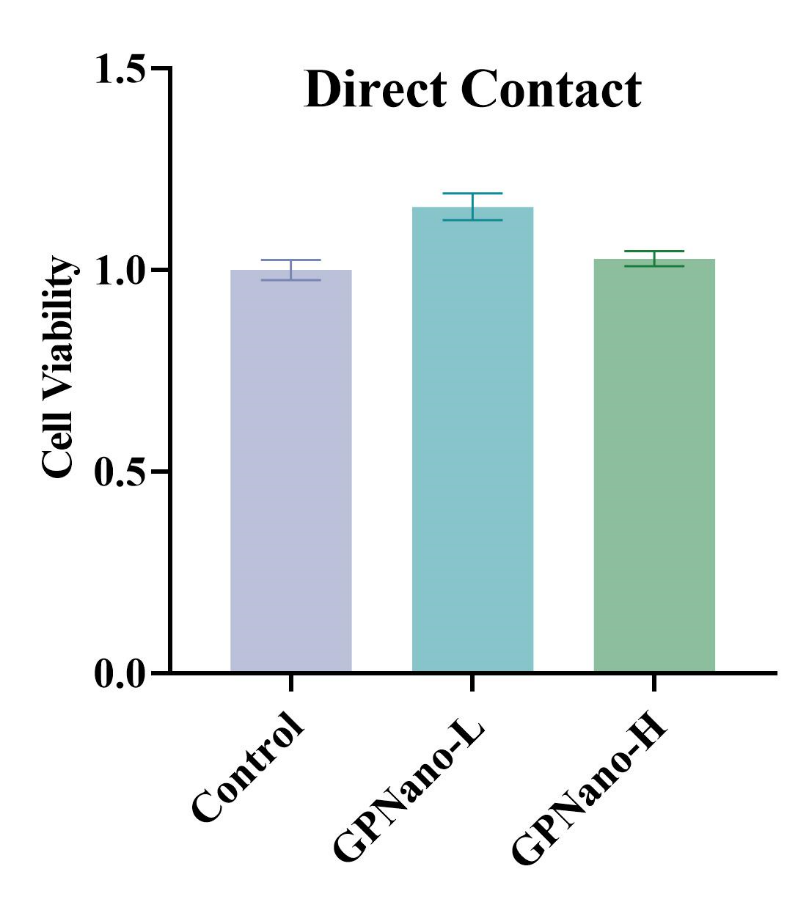

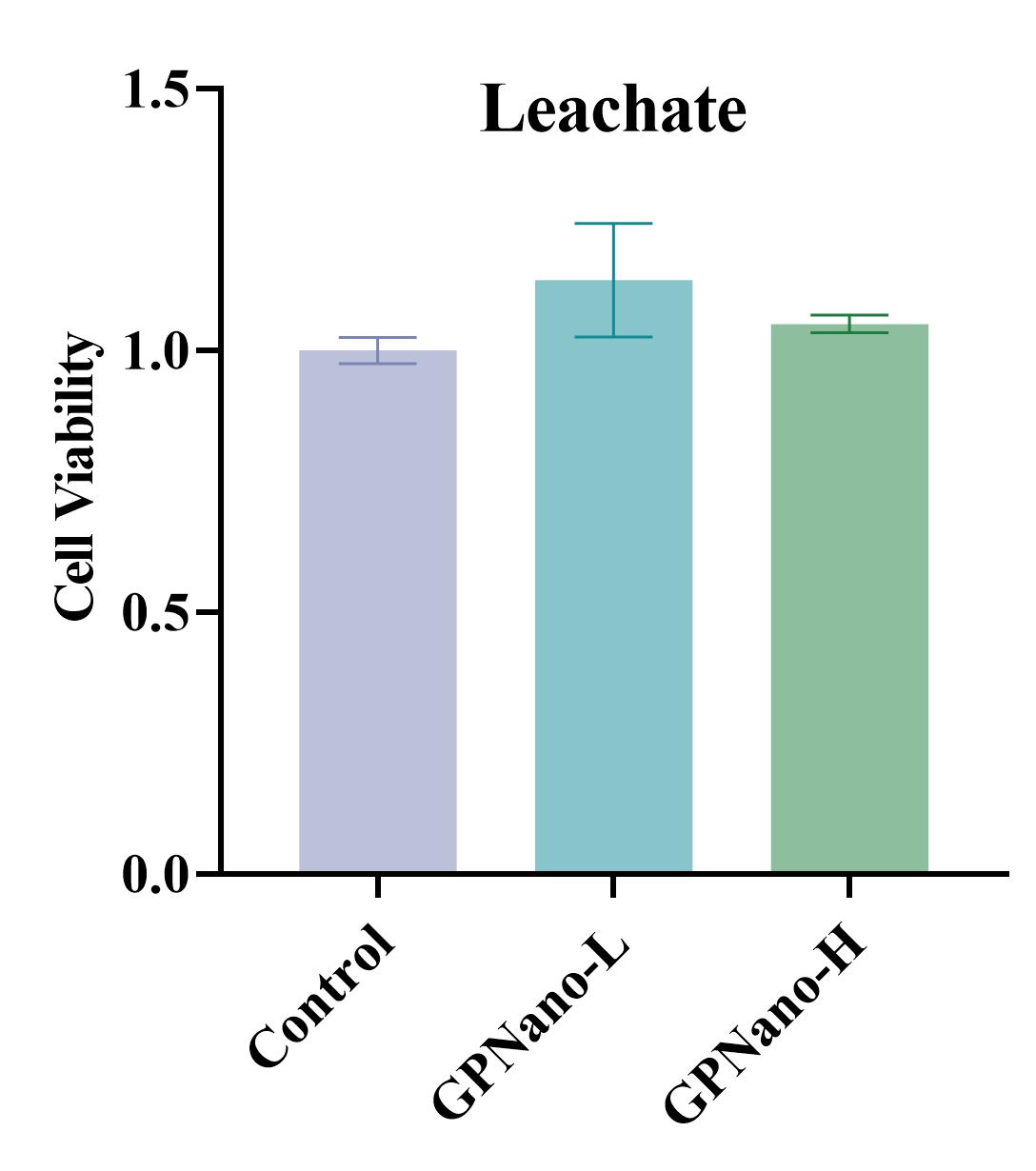

**Supplementary Fig. 12** Quantifications of different cell viability tests measured by CCK-8 (n=3, error bar represents the standard error of means).

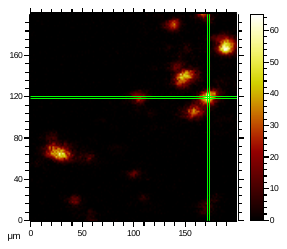

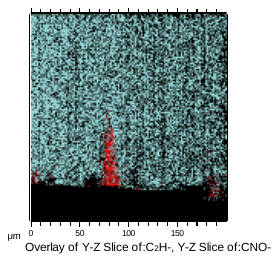

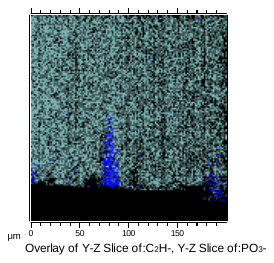

Accumulated CNO^-^ X-Y map

Y-Z cross-section.

C_2_H^-^ overlayed with PO_3_^-^

Thickness of GPNano-H

Thickness of GPNano-H

200 μm

200 μm

200 μm

200 μm

**Supplementary Fig. 13** 2D cross-sectional analysis of ToF-SIMS negative ions. Pale blue: C_2_H^-^. Red: CNO^-^. Blue: PO_3_^-^

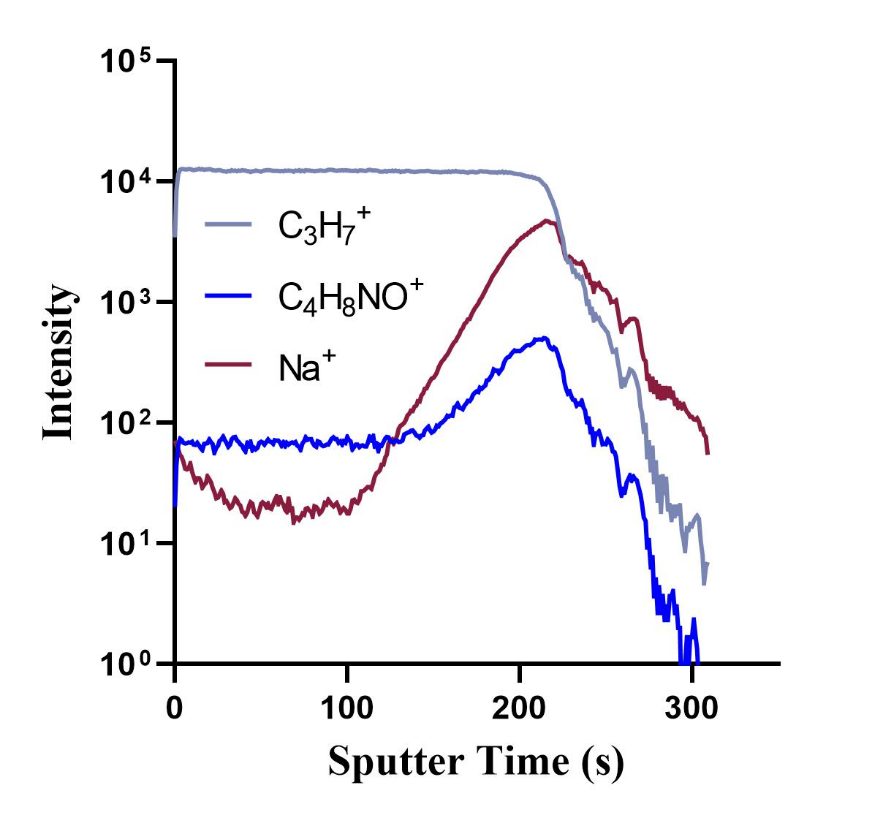

**Supplementary Fig. 14** Positive ToF-SIMS depth profile analysis of GPNano-H scanning from the air site to the wound site

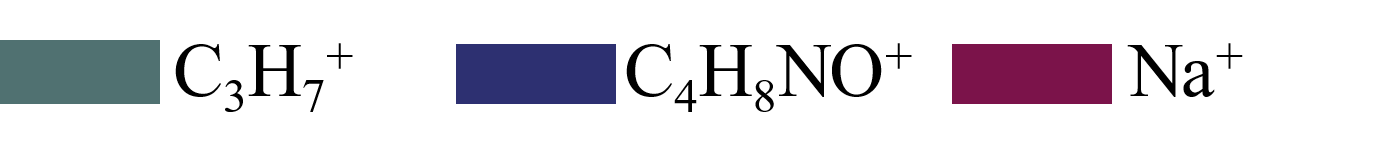

200 μm

200 μm

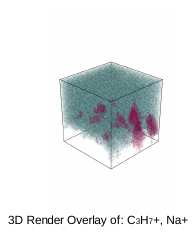

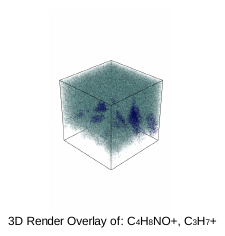

Thickness of GPNano-H

**Supplementary Fig. 15** 3D ToF-SIMS rendering analysis of GPNano-H where C_3_H_7_^+^ representing the UHMWPE is overlayed with C_6_H_8_NO^+^ and Na^+^ representing the wound exudate-related species.

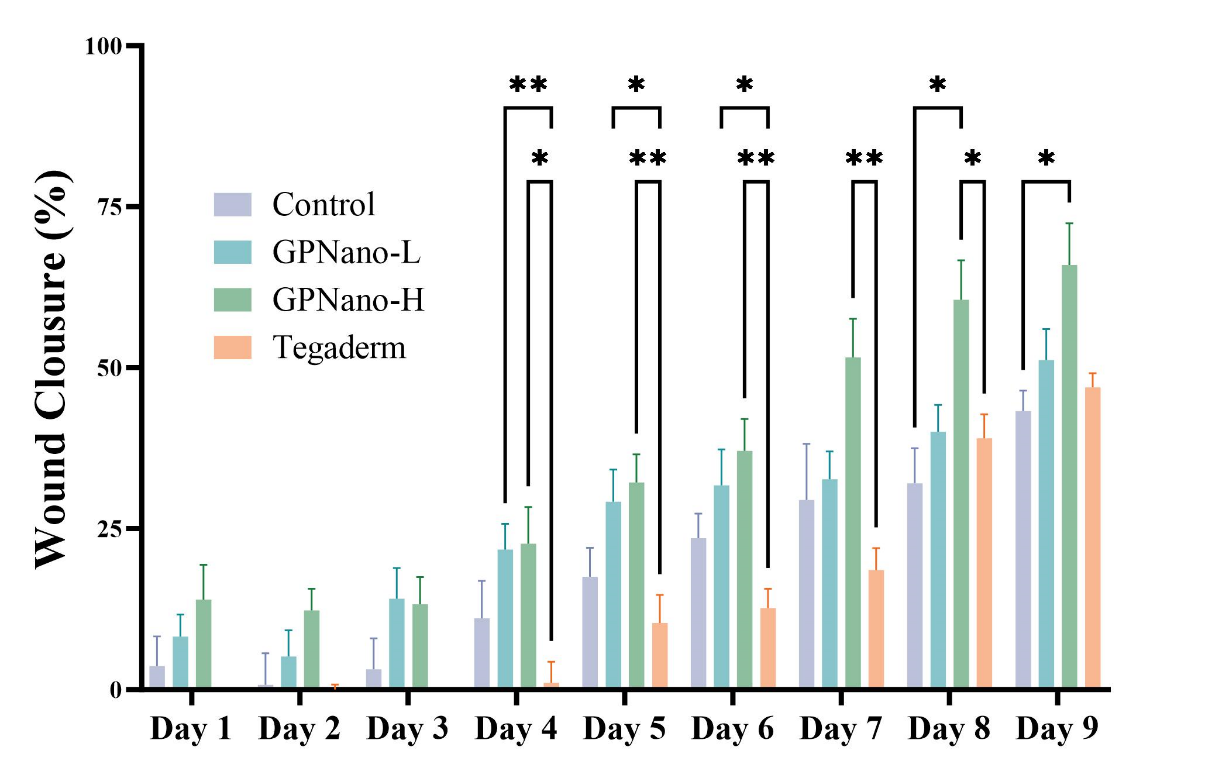
**Supplementary Fig. 16** Relative wound closure rate from day 1 to day 9 (n=5, *p<0.05, **p<0.001, error bars represent the standard error of means).

**Supplementary Fig. 1**
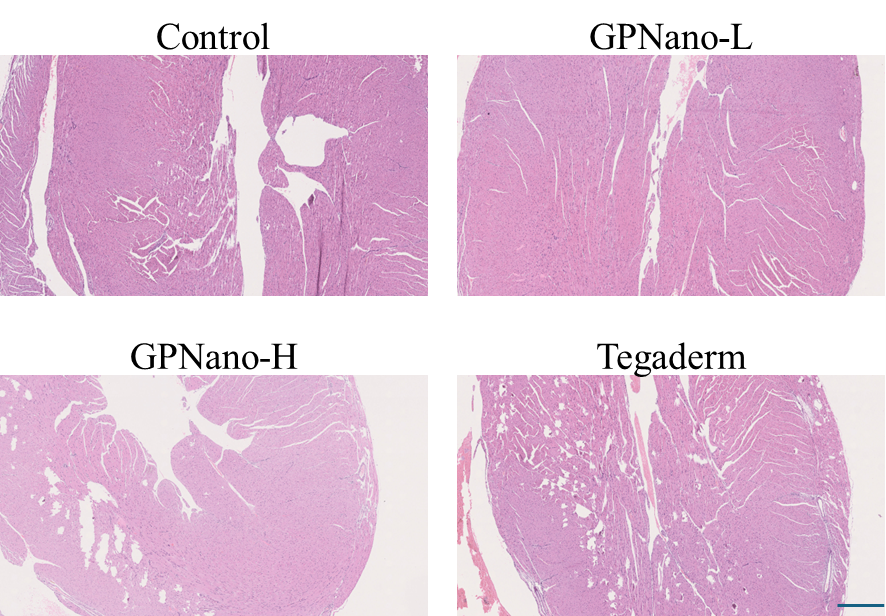
**7** Representative images of H&E-stained heart tissue. Scale bar: 500 μm.

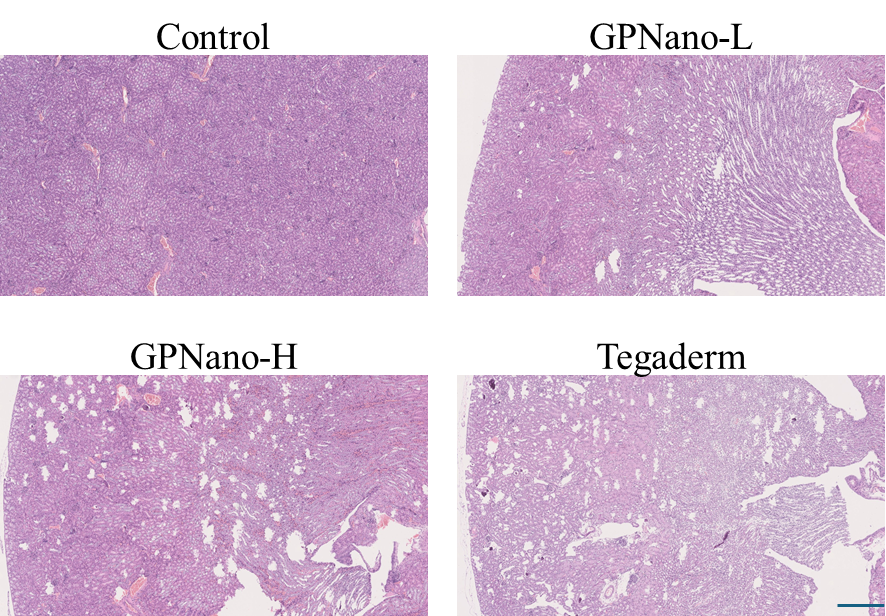
**Supplementary Fig. 18** Representative images of H&E-stained kidney tissue. Scale bar: 500 μm.

**Supplementary Fig. 19** Representative images of H&E-stained liver tissue. Scale bar: 500 μ
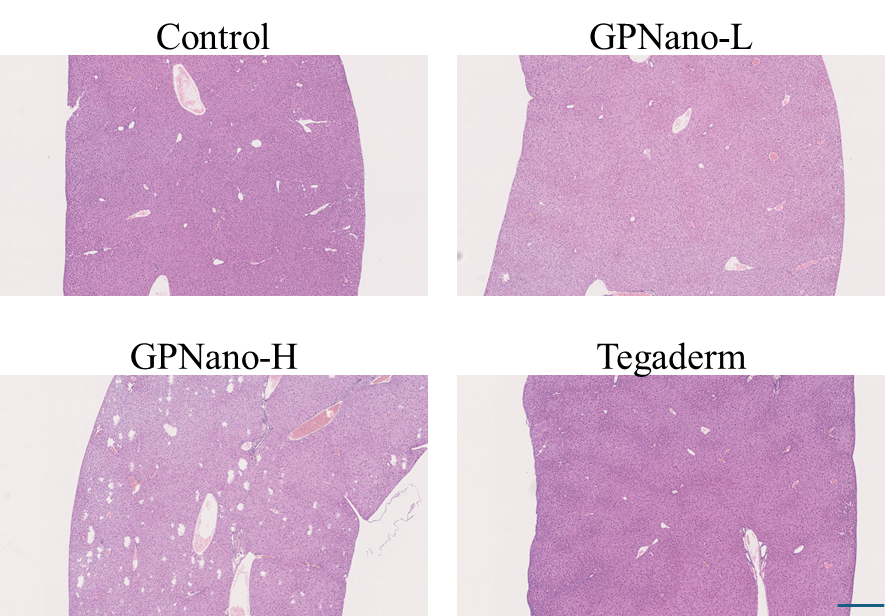
m.

**Supplementary Fig. 20** Representative images of H&E-stained spleen tissue. Scale bar: 500 μ
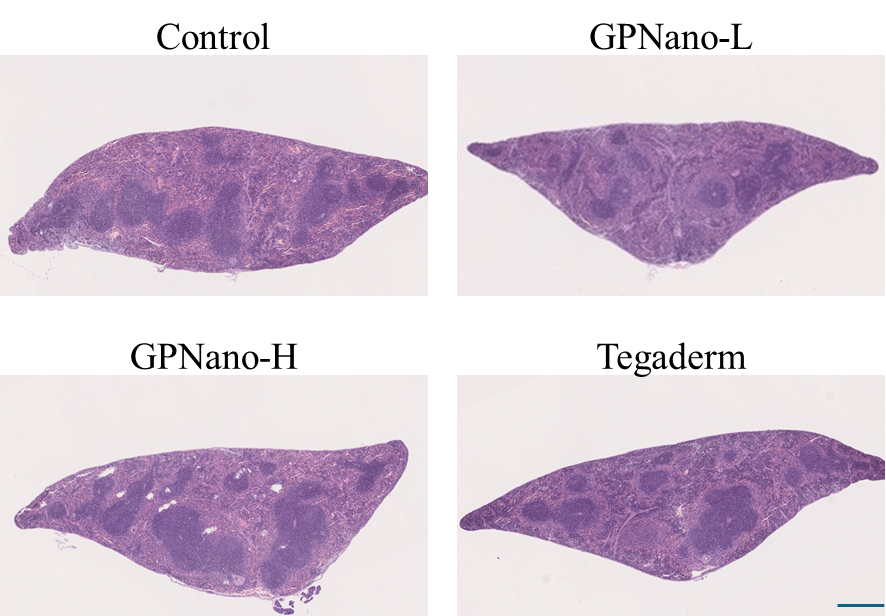
m.

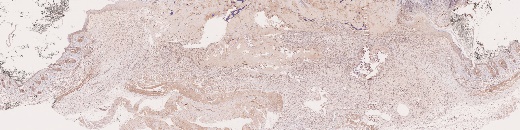

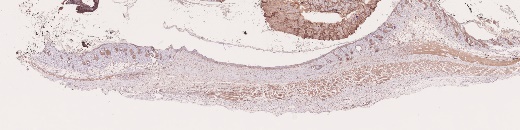

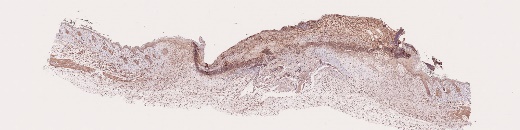

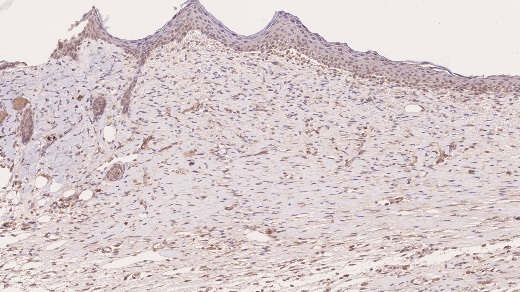

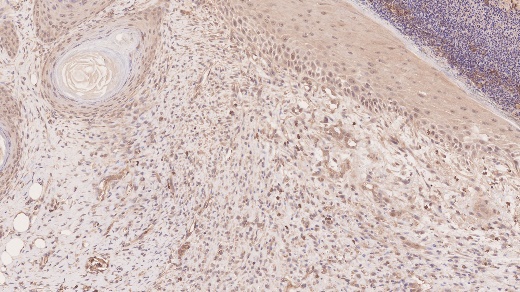

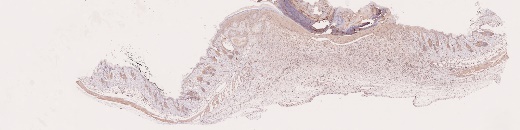

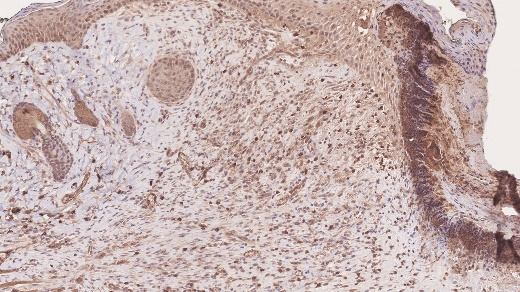

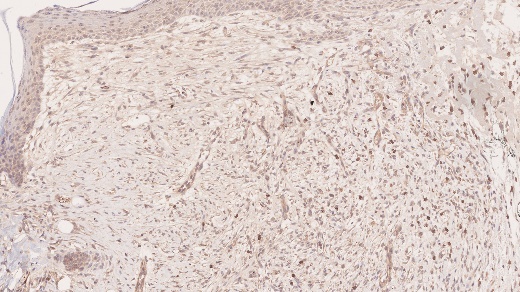

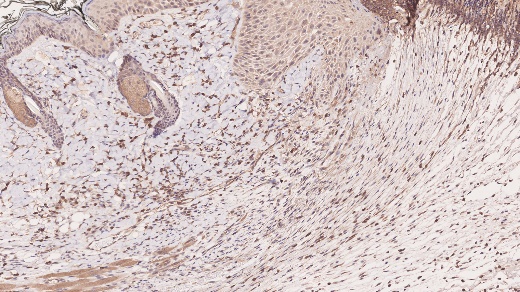

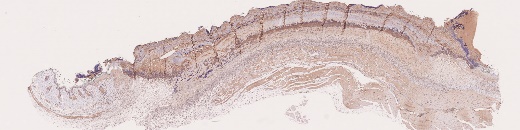

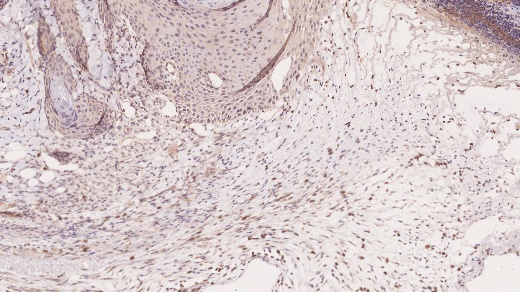

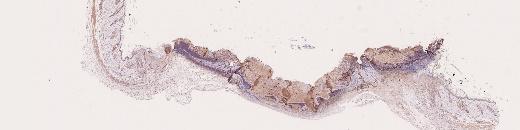

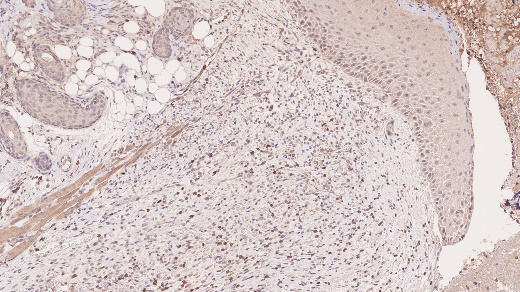

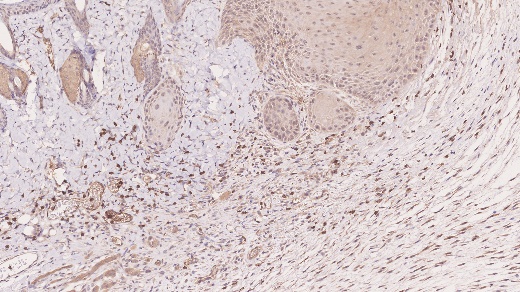

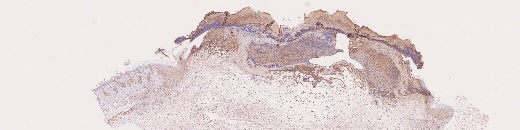

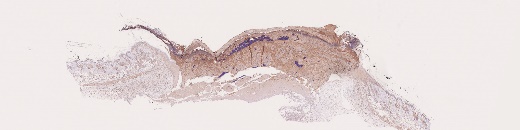

Day 5

Day 9

Control

GPNano-L

GPNano-H

Tegaderm

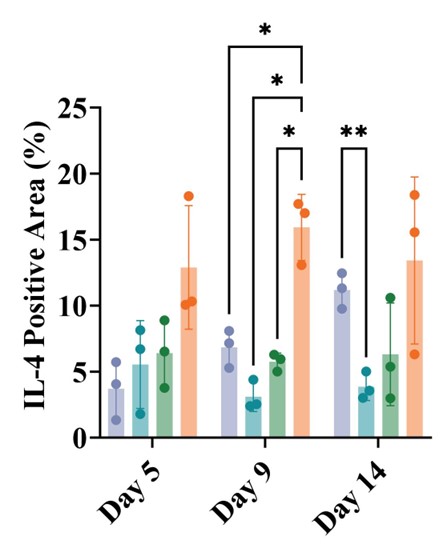

**Supplementary Fig. 21** Representative immunohistochemistry staining of IL-4 on day 5 and day 5 (Brown: IL-13. magnification, x25, scale bar: 1 mm; magnification, x200, scale bar: 300 μm) and quantification analysis of IL-4 expression from day 5 to day 14. (n=3, one-way ANOVA. *p<0.05, **p<0.01, error bars represent the standard error of means).

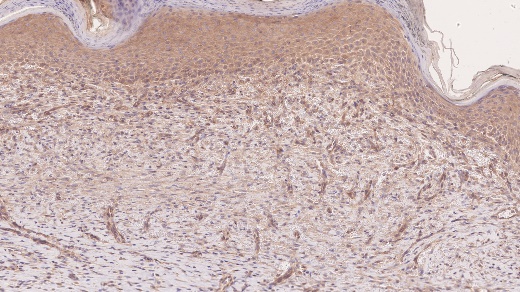

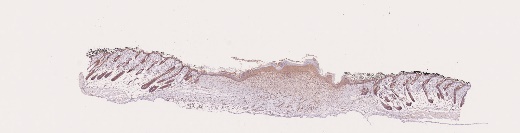

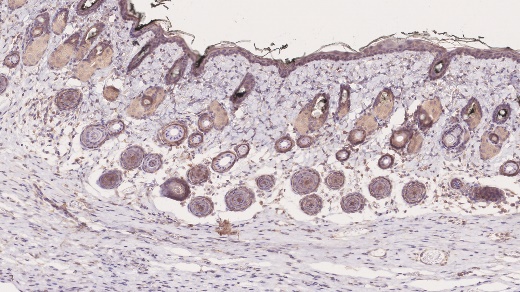

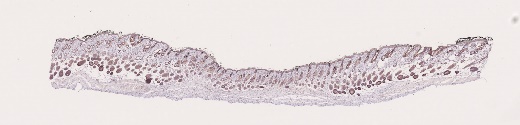

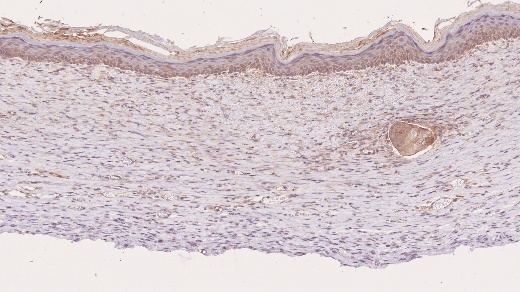

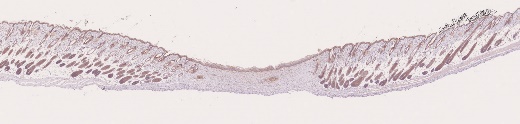

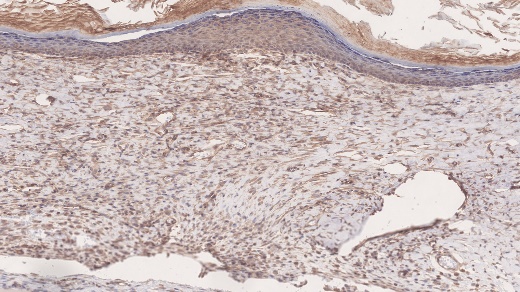

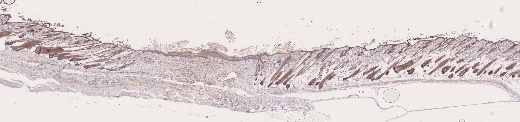

Day 14

Control

GPNano-L

GPNano-H

Tegaderm

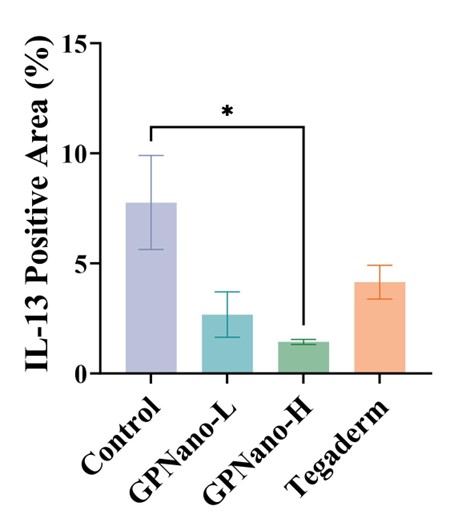

**Supplementary Fig. 22** Representative immunohistochemistry staining of IL-13 on day 14. (Brown: IL-13. magnification, x25, scale bar: 1 mm; magnification, x200, scale bar: 300 μm) and quantification analysis of IL-13 expression on day 14. (n=3, one-way ANOVA. *p<0.05, error bars represent the standard error of means).

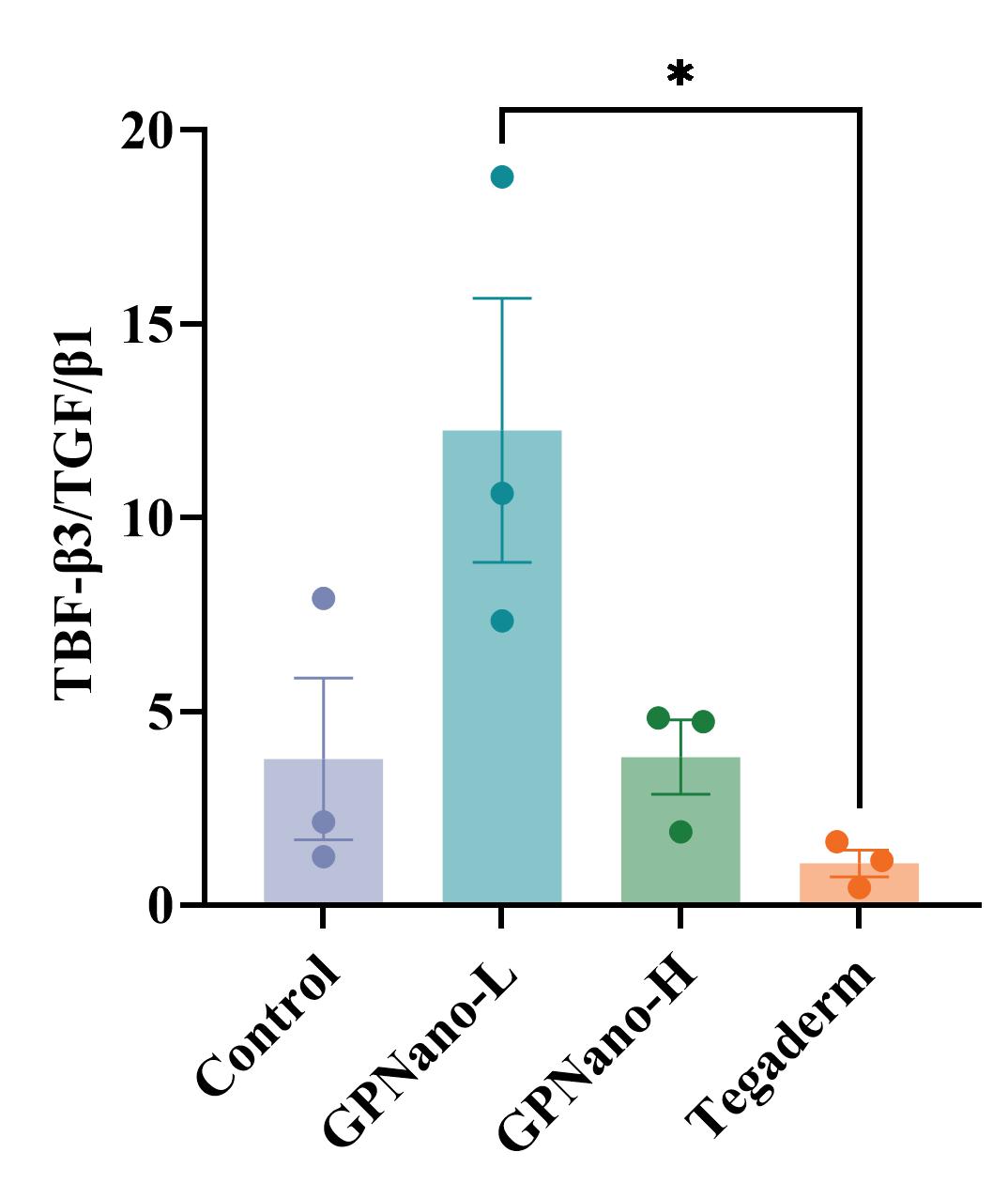
**Supplementary Fig. 23** Quantification of the TGFβ3/TGFβ1 ratio. (n=3, unpaired t-test, *p<0.05, error bars represent the standard error of means).

**Supplementary Fig. 24** Schematic illustration of how to set up a peel strength test on utPE samples.

Sample

Tape 1

Tape 2

**Supplementary Fig. 2**
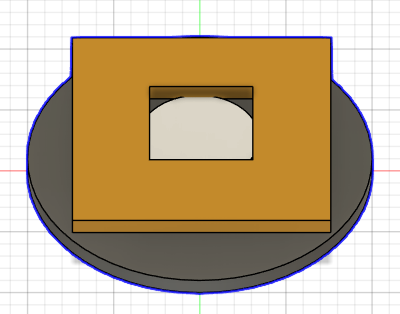
**5** Schematic illustration of how to set up a ToF-SIMS sample. Orange: SEM tape; white: utPE sample; black: silicon wafer.
